# Catch-slip behavior observed upon rupturing membrane-cytoskeleton bonds

**DOI:** 10.1101/184069

**Authors:** Vivek Rajasekharan, Varun K. A. Sreenivasan, Fred A. Pereira, Brenda Farrell

**Affiliations:** Department of Otolaryngology – Head and Neck Surgery, Baylor College of Medicine, Houston, TX 77030; Department of Molecular and Cellular Biology, Baylor College of Medicine, Houston, TX 77030; Current affiliation: Lowy Cancer Research Centre, Level 3, University of New South Wales, Sydney, NSW 2052, Australia; Huffington Center on Aging, Baylor College of Medicine, Houston, TX 77030

## Abstract

Cells are capable of cytoskeleton remodeling in response to environmental cues at the plasma membrane. The propensity to remodel in response to a mechanical stimulus is reflected in part by the lifetime of the membrane-cytoskeleton bonds upon application of a tensile loading rate. We measure the lifetime and force to rupture membrane-cytoskeleton linkages of a head and neck squamous cell carcinoma (HNSCC) cell line, HN-31 by applying a tensile loading rate (< 60 pN/s) with a handle bound to a cell, while monitoring the displacement of the handle at 2 kHz after averaging. We observe the lifetime increases with loading rate, *r_f_* to a maximum after which it decreases with further increase in *r_f_*. This biphasic relationship appears insensitive to drugs that target microtubule assembly, but is no longer detectable, i.e., lifetime is independent of *r_f_* in cells with reduced active Rho-GTPases. The loading rate-time relationship resembles catch-slip behavior reported upon applying tensile loads to separate protein complexes. Under small loads the bonds catch to increase lifetimes, under larger loads their lifetime shortens and they dissociate in a slip-like manner. Our data conforms to a model that considers the membrane-cytoskeleton bonds exhibit a load-dependent conformational change and dissociate via two pathways. We also find the membrane-cytoskeleton linkages strengthen with stationary compressive load, *F_SC_* (|*F_SC_*| < 40 pN), and conclude this metastatic cell line responds to small mechanical stimuli by promoting cytoskeleton remodeling as evident by observing F-actin within the membrane nanotube (10 µm length) formed after bond rupture.

## 1. Introduction

Non-covalent bonds between two binding proteins break and form stochastically (energy per bond < ∼10 *κ_B_T*^1^), and this dynamic behavior is considered crucial for their biological activity. The rate at which these bonds form and break depends upon the magnitude of the activation energy barrier^2, 3^ that separates the bonded (A-B) from the un-bonded state^4^ (A, B) and is affected by temperature, viscosity of the medium, and external force. These theories were initially developed to describe the kinetics of chemical reactions occurring within gases or liquids and were later applied to model dissociation of proteins *in vitro* or *in situ*^5^ (i.e., within the cytoplasm of cells or between cells). Bell modelled bond rupture as cells are pulled apart by an applied tensile force where it was predicted that the applied tensile force should increase the off-rate, *k* (shortens the lifetime of bonds) by reducing the activation energy barrier, i.e., 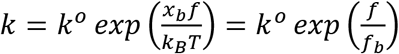 where *κ*_*B*_ is the Boltzmann constant, *T* is the absolute temperature, *kº* is the unstressed off-rate, *f* is the applied force, *x*_*b*_ is the energy barrier distance and 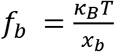 is the thermal force scale for a single bond and 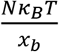 for *N* bonds^5^ (refer to Table S1 for list of all abbreviations). Indeed, experiments performed to break a range of non-covalent bonds with different techniques, flow chamber^6^, atomic force microscopy^7^ (AFM), optical tweezers^8^ (OT), and biomembrane force probe^9^ (BFP) show that application of a tensile force reduces the bond strength and the lifetime of the bonds in agreement with the model. This model predicts that an applied compressive force should increase the lifetime of the bonds by increasing the on-rates. This was demonstrated for interactions of integrin with intercellular adhesion molecule 1 (ICAM-1)^10^ and L-selectin or E-selectin with P-selectin glycoprotein ligand 1 (PSGL-1)^11, 12^.

In contrast to the intuitive Bell model for how load affects the lifetime of bonds, Dembo theorized that certain bonds may be strengthened by application of a tensile load to exhibit lifetimes that increase with a tensile load. This non-intuitive conjecture led him to coin them *catch* bonds^13^. Experimental evidence for the existence of catch bonds was demonstrated when *Escherichia Coli* (*E. Coli*) were found to be more firmly attached to red blood cells (RBCs) at high shear stress. In this example, the receptor on *E. Coli* is an adhesin (FimH) and the ligand on the RBC is a sugar (i.e., mannose) of a glycoprotein^14^. Further experimental evidence for catch behavior was reported for the bond between another adhesion molecule P-selectin and its ligand, PSGL-1 in single molecule *in vitro* studies with AFM^15^ when used in force clamp mode. They found these bonds stabilize and have prolonged lifetimes with increasing load, i.e., they are catch-like until a threshold after which they destabilize and exhibit shortened lifetimes, i.e., become slip-like. They coined this transition *catch-slip*, where maximal lifetime is measured at the catch-slip transition and is 1.1 s for P-selectin and PSGL-1^15^. Similar single molecule experiments performed between FimH and its ligand, mannose also show the catch-slip transition with a longer maximum lifetime of ∼25 s^16^. More recently catch behavior was observed when breaking bonds between other cell-adhesion proteins and their ligands. They were detected between integrin *α_5_β_1_* and fibronectin with AFM^17^ and between integrin *α_L_β_2_* expressed on a neutrophil and ICAM-1 with a BFP^18^. The maximal lifetimes of these bonds were 1-3 s. Catch bonds were also observed upon separating two monomeric E-cadherins that make up a dimer where the maximal lifetime of this bond was much less at 70 ms^19^.

Cytoskeletal motor proteins are also found to exhibit catch-like behavior^20–23^. Catch bonds were detected between F-actin and myosin II^20^ and between F-actin and myosin 1b^21^. In both cases their maximum lifetimes are up to 25 ms. Catch-slip behavior was also detected upon separating monomeric G-actin from an F-actin filament to exhibit a maximum lifetime of 1 s^22^. A catch bond with a significantly longer lifetime of 50 minutes was reported between kinetochore and microtubules^23^. These observations of catch bonds in cytoskeletal motor proteins and cell surface receptors reveal a common theme that the functional role of these bonds is to support a load or transmit a force^24^.

Although most observations of catch bonds are with reconstituted purified systems *in vitro*, Chen et al. detected catch bonds on a living cell; they formed bonds between extracellular domains of integrins expressed on the membrane of Jurkat cells and their ligands (expressed on the BFP probe) and observed catch behavior upon application of a tensile force on these bonds^18^. Despite evidence that cytoskeletal proteins (e.g., F-actin^22^, myosin II^20^, myosin 1b^21^, and microtubules^23^) can form catch bonds, there are no reports demonstrating catch behavior within the cytoskeleton assemblies of a living cell. One report measured the force to rupture the membrane-cytoskeleton bonds of neutrophils under very high tensile loading rate (250 to 38,000 pN/s) with BFP, but only found evidence for Bell-like bonds^25^.

We utilize a similar technique^26^ to determine the lifetime of membrane-cytoskeleton bonds at the edge of a cancer cell. This technique was first developed to estimate the non-local bending moduli of membranes of vesicles^27^ and plasma membranes of RBCs^28^. The technique was adapted to determine membrane tension and bending moduli of plasma membranes of other cells (e.g., neurons^29^, neutrophils^30^) by measuring the force to form a membrane nanotube or tether from the cell. These studies suggest that the major contribution to forming a nanotube from cells was overcoming adhesion energy – i.e., breaking the bonds between the membrane and cytoskeleton. In these studies, the force was reported at slow rates^29, 31–34^ (< 100 Hz) and the adhesion energies were estimated from this force. The time course of this force was not reported making it difficult to infer kinetics of the membrane-cytoskeleton bonds. In the presented work, we break the plasma membrane-cytoskeleton bonds of a cell and monitor both the force and the time course of the force with a resolution limited by the corner frequency of our instrument at 2.1 ms^26^. The experiments are performed with HN-31 cancer cells: a head and neck squamous cell carcinoma (HNSCC) cell line^35^ that exhibits a wide range of mutations^36^ including mutated *TP53*^37^ and an *H-ras* mutation^38^ that results in persistently-active Rho-GTPases. We apply a constant loading rate to rupture the bonds and detect a biphasic time-loading rate relationship. This biphasic relationship resembles catch-slip behavior suggesting that membrane-cytoskeleton bonds in these cells can stabilize and increase their lifetimes up to a threshold loading rate. Various models^39–41^ attribute this catching behavior to a load-dependent conformational change that precedes the dissociation event; we adapt a model^39^ to describe this biphasic time-loading rate relationship.

## 2. Materials and Methods

### 1. Cell Culture

HN-31, a metastatic HNSCC cell line is maintained by culturing in Dulbecco’s modified Eagle’s medium (Gibco/Invitrogen, Grand Island, NY) with the following additives: 10% (v/v) fetal bovine serum, 250 U/ml penicillin – 250 µg/ml streptomycin, minimal essential medium vitamin solution, sodium pyruvate (Thermo Fisher Scientific, Waltham, MA), and non-essential amino acids and L-glutamine (Lonza Group Ltd., Basel, Switzerland). Cells are incubated at 37ºC with 5% CO_2_ atmosphere and grown to ∼80 % confluence. For each experiment, cells are re-seeded onto a 35 mm poly-D-lysine-coated glass bottom Petri dish (MaTek Corp., Ashland, MA) and grown to < 30% confluence.

### 2. Physiological Saline Solution

Buffered saline contains (in mM) 150 NaCl, 2 CaCl_2_, 1 MgCl_2_, 10 HEPES and 2.8 KOH. The pH and osmolality of the solution are adjusted to 7.2 and 300 mOsm Kg^-1^ by adding NaOH and glucose, respectively. All force measurements are performed and completed within an hour of replacing the culture media with buffered saline.

### 3. Drug Treatments

Nocodazole (EMD Chemicals, Inc., San Diego, CA) dissolved in DMSO (Calbiochem, San Diego, CA) is added at a final concentration of 15 µM to disrupt microtubules^42^. Latrunculin A (AdipoGen, San Diego, CA) dissolved in DMSO is added at a final concentration of 1.7 µM to inhibit the polymerization of actin^43^. Both drugs or the solvent DMSO are added to the buffered saline within the Petri dish at least 20 minutes prior to an experiment, and remain in the solution for the duration of an experiment, i.e., 40 min. *C. difficile* Toxin B (List Biological Laboratories, Inc., Campbell, CA) at a concentration 200 ng/ml was used to non-specifically inhibit Rho-GTPases. As inhibition of Rho-GTPases with Toxin B is not reversible^44^ cells were incubated with the drug in serum-free media for 2 hours for the pull-down assay and force measurements or they were incubated for 4 hours for pull-down assay. The force measurements were then performed in buffered saline.

### 4. Fluorescence Microscopy

We visualized F-actin by fluorescent labelling in untreated cells, cells incubated with Latrunculin A for 20, 40 and 60 minutes in buffered saline, and cells incubated in serum-free media with Toxin B for 2 and 4 hours. Cells are (all percentages are volume/volume): (i) rinsed with buffered saline; (ii) fixed with 4% paraformaldehyde (Electron Microscopy Sciences, Hatfield, PA) in buffered saline for 10 min; (iii) rinsed with buffered saline; (iv) permeabilized with 0.1 % triton X-100 (Sigma-Aldrich, St. Louis, MO) for 5 min; (v) rinsed with buffered saline; (vi) incubated with 2.5 % bovine serum albumin (Sigma-Aldrich) for 30 minutes to reduce non-specific binding; and (vii) stained with phalloidin (Alexa Fluor 488, Thermo Fisher Scientific) at 37 ºC for 1 hour; and (viii) rinsed with buffered saline. To image membrane nanotubes formed from cells, the bead used to form the nanotube is adhered to the Petri dish by moving the Petri dish upwards towards the bead (in the *+z* direction) with the 3-D piezo stage until contact. We then fix and stain for F-actin with Alexa 488 phalloidin (steps (ii) to (viii)) as described above. To visualize microtubules, untreated cells and cells incubated with Nocodazole for 20, 40 and 60 minutes in buffered saline are treated as described in steps (i) to (vi). Then the cells are incubated with the primary antibody, anti-tubulin polyclonal antibody (ATN02, Cytoskeleton, Inc., Denver, CO) for 1 hour, washed thrice and incubated with the secondary antibody, anti-sheep Alexa Fluor 488 (Thermo Fisher Scientific) for 30 minutes and rinsed. All cells are visualized with a Leica DM2500 confocal fluorescence microscope (Leica Microsystems, Wetzlar, Germany) and images are captured with the Leica LAS AF 2.6.0 software. For each treatment and the untreated cells, the concentration of staining agent added and the illumination and detection conditions (e.g., laser power, exposure time) were constant.

### 5. RhoA GTPase Pull-down Assay

Reduction in RhoA-GTPase activity is assessed after 2 and 4 hours of treatment with Toxin B with the RhoA Pull-down Activation Assay Biochem Kit (Cytoskeleton, Inc.). The assay utilizes agarose beads bound to effector domain Rhotekin-RBD to pull-down the active GTP-bound form of RhoA-GTPase. We follow the protocol described by the manufacturer to determine reduction in RhoA-GTPase activity after drug treatment. Protein samples were analyzed by SDS-PAGE and western blot with primary antibodies provided with the kit. β-tubulin is used to normalize both total and active RhoA band intensities to account for differences in protein loading [e.g., Normalized RhoA = 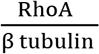]. Active RhoA intensity: total RhoA intensity (RhoA Ratio) is determined for each treatment 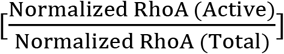. Reduced activity after addition of drug for each time point is determined with respect to untreated cells [e.g., 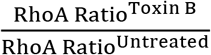]. All ratiometric fluorescence changes were quantified with Odyssey imaging system (LI-COR, Lincoln, NE). Note the antibody provided with the kit resulted in low intensity Rac1 bands, so no further experiments were performed with Rac1. The antibody for Cdc42 resulted in a band approximately triple the weight, which could be a trimer or non-specific binding (spiking the loading buffer with fresh β-mercaptoethanol and boiling the sample did not change the outcome). Therefore, the change in the activity of these Rho-GTPases with Toxin B treatment was not measured.

### 6. Optical Tweezers

A custom built optical tweezers is used. The instrumentation, calibration and noise determination have been reported in detail^26^. We only describe the main features. The optical tweezers is formed with a continuous wave Ti-sapphire laser (3900S, Spectra-Physics, Santa Clara, CA) tuned to 830 nm and pumped by a 532 nm solid-state frequency-doubled Nd: YVO_4_ laser (Millennia Prime, Spectra-Physics) with a maximum power of 6 W. The beam is then steered via an optical path to fill the back aperture of an objective lens (numerical aperture of 1.4) within an inverted microscope (Eclipse-Ti, Nikon Instruments Inc., Melville, NY). The laser power and position of the beam are controlled by two acousto-optic deflectors, AODs (72003 and R21.35-65-2ASVCO-2, Gooch & Housego, Melbourne, FL). The beam position is steered with a field programmable gate array, FPGA (PCIe-7852R, National Instruments, Austin TX) with LabVIEW (from v.12, National Instruments). A fluorescent bead (FluoSpheres Sulfate Microspheres, F-8858, Life Technologies, Carlsbad, CA) with excitation maxima 576 nm, emission maxima 605 nm and radius 2 µm is trapped. The bead fluorescence is excited with a TLED (TLED+, Sutter Instrument Company, Novato, LA) or xenon lamp (Lambda LS xenon arc, Sutter Instrument Company) and the emission is filtered with a bandpass filter. The position *(X, Y)* and intensity of the emitted light are detected by a fast quadrant photodiode, QPD (2901, New Focus, Newport, NY) with linear range ± 2 µm. The QPD voltages are acquired with a digital acquisition card, DAQ (PCIe-6363, National Instruments) at 200 kHz with LabVIEW.

The instrument is calibrated with a variation of the drag force method^26^. At the laser power employed for the experiments (0.130 W at objective), the typical corner frequency is ∼470 Hz which results in a stiffness *K* of 0.116 pN/nm (±0.011 pN/nm) at the height of the trapped bead above the dish (9.88 ± 0.04 µm), the radius of bead (2.13 ± 0.02 µm) and the viscosity of buffered saline (0.920 mNsm^-2^).

### 7. Data conditioning and analysis

The voltage outputs of the QPD are *X, Y* and the sum (Σ); *X* and *Y* carry positional information of the bead and the third channel, the Σ contains the intensity information. The signal is averaged hundred-fold post collection to result in a sampling frequency of 2 kHz. Further processing is performed with custom written MATLAB (from vR2011b, The Mathworks, Natick, MA) scripts. We check the Σ to find whether there are artifacts in the signal due to floating beads entering the illuminated area, when found this data is discarded. To calculate the displacement of the bead *(x,y)* we translate the voltage output from the QPD to nm. This requires executing three steps. (i) The Σ decreases with time due to photo-bleaching of the bead. To correct for photo-bleaching we fit the sum to a polynomial of order two and divide the *X* and the *Y* signals with this quadratic fit to the Σ. (ii) The method we use to convert these displacements from normalized voltage in V/V to nm has been described^26^. Briefly, the trapped bead is displaced with the AODs predetermined distances (i.e., −800, −500, +500 and +800 nm) in the *x* direction, and the output from the QPD is measured for each distance. We then determine the linear slope *m_QPDX_* (V/V/nm) between the bead displacement and normalized voltage. (iii) To determine the displacement (*Δx*) in nm during an experiment we divide the normalized voltage signal (V/V) determined in (i) with *m_QPDX_*. We then plot *Δx* versus time for the first 10 s of the experiment when no force is acting on the bead (i.e., *Δx* should be zero). A small offset is usually observed due to the initial position of the bead being slightly off center of the QPD. To correct the offset, we determine the mean *Δx* for the first 5 s and subtract it from *Δx* for the entire recording. The force on the bead is the product of final *Δx* and *K*. We determine the noise in the data for each experiment by fitting the position vector (*r_m_, θ*) to a Rayleigh (*r_m_*) and uniform (*θ*) distribution as described^26^. The scale factor of the Rayleigh distribution is ∼22 nm (up to 32 nm with one TLED lamp) at 2 kHz. Deviation of the angle from the uniform distribution indicates the presence of an external force. Uniform distribution is observed when the bead separates from the plasma membrane and can move in all directions due to the absence of force.

## 3. Results and Discussion

### 1. Determination of the lifetime, force to rupture and loading rate

At the beginning of the experiment, the cell attached to a dish is placed approximately 20 µm apart from an optically trapped bead as shown in Figure 1A and monitoring of the position of the trapped bead commences. After recording for 10 s, the cell is moved towards the trapped bead in the *x* direction with a 3-D piezo stage (Physik Instrumente L.P., Auburn, MA) at velocity *v*: 0.5 µm/s for 40 s. In some cases, to prevent very large attractive forces upon cell bead contact, that could push the bead out of the trap, the 3-D piezo stage was stopped 1-2 s earlier. Typically at this time (−12 to −10 s), the bodies interact and the bead exerts a compressive load (negative, magnitude up to 40 pN) on the cell (Figure 1B); for the example shown, the bodies interact at 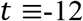 s and the cell is pushed into the bead for a further 2 s with the force reaching −30 pN when *v* is terminated (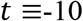 s, Figure 1D). The cell and bead are left in contact for 10 s to ensure adhesion during which the force relaxes reaching a stationary compressive load, *F*_*sc*_ of −18 pN at 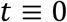 s. The cell is then pulled away from the bead in the *x* direction at the same velocity (Figure 1C and D) for 20 s. The membrane-cytoskeleton bonds rupture after which a membrane nanotube forms (Figure 1D Inset). Note: we probe each cell with a bead only once; we do not repeat the experiment on the same cell even in cases where no nanotube has formed since the loading history can influence bond rupture^20, 40, 45^.

**Figure 1:**
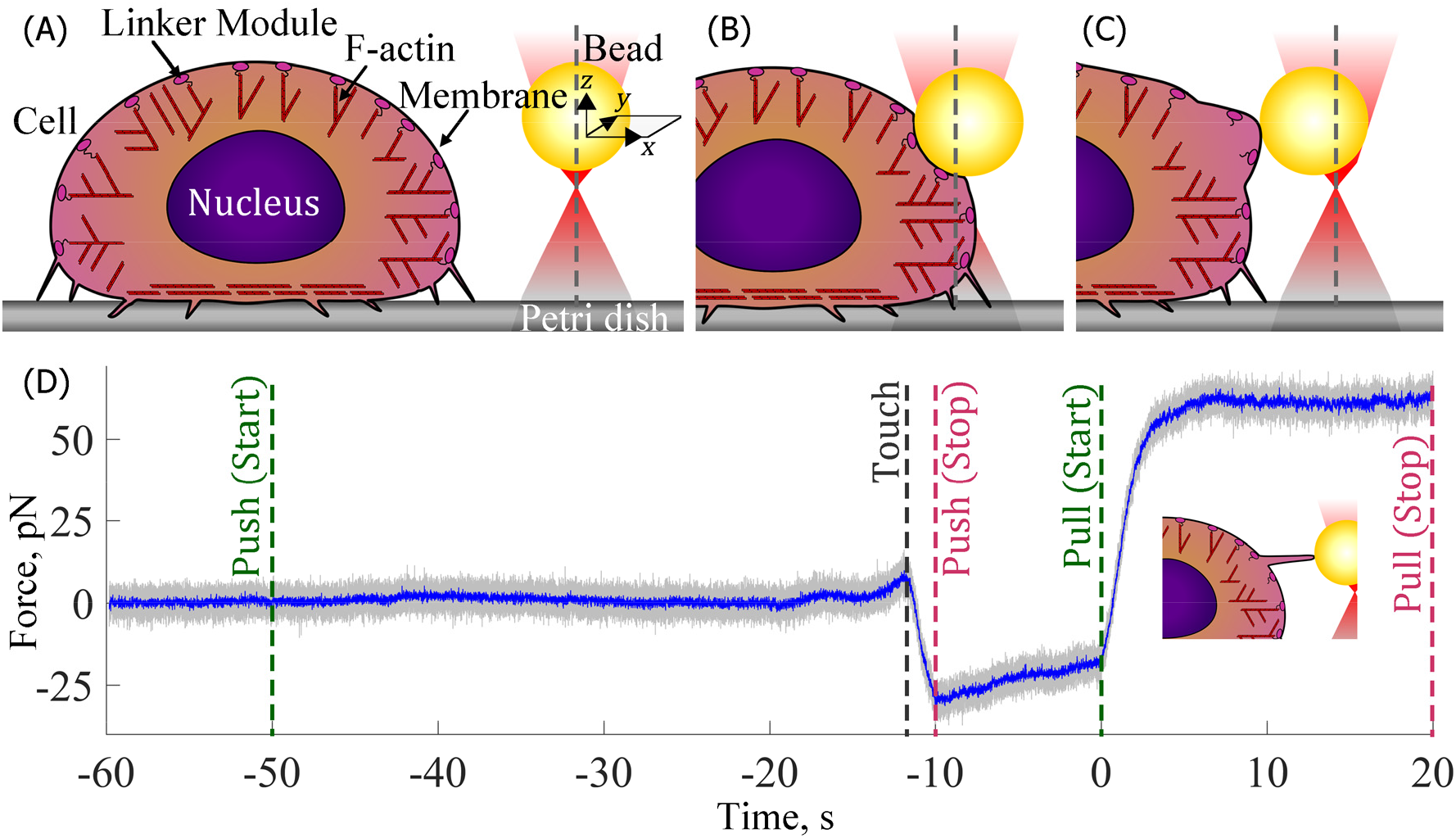
Description of an experiment. The measured force with respect to time is shown in D with schematics of the process shown in A, B and C. At the onset of the experiment a cell attached to a Petri dish and a trapped bead *(K:* 0.116 pN/nm) are 20 µm apart and held in that position for 10 s (**A & D**). At time indicated by *Push (Start)*, the cell is brought towards the bead with a 3-D piezo stage (*v*: 0.5 µm/s) in the *x* direction. The bodies touch (at time indicated by *Touch*) and interact under the compressive loading rate till the time indicated by *Push (Stop)* after which motion is stopped to enable the bodies to adhere (**B & D**). To rupture the membrane-cytoskeleton bonds the cell is pulled away (*x* direction) from the bead starting at time *zero* indicated by *Pull (Start)* through time indicated by *Pull (Stop)* (**C & D**). Depending upon the strength of membrane-cytoskeleton bonds the bead moves in the same *x* direction towards the cell at a measured loading rate that is detected as a linear rise in force. Upon rupture of the membrane-cytoskeleton bonds a membrane nanotube forms (**D, Inset**) that extends easily to a length of ∼10 µm, the maximum pulling distance. In this example, the force plateaus as there is no further resistance to nanotube elongation i.e., membrane tension is constant with time. The sampling frequency of the force is 2000 (*grey*) and 200 Hz (*blue*)

Upon application of a tensile load the force increases linearly with time until a critical force, *f*^*R*^ is attained, which is detected by a significant drop in the slope. This change in slope indicates membrane-cytoskeleton bond rupture, and was evident by detecting one of the three scenarios (Figure 2, Top and Middle Panels): (i) the force drops to a constant value and the slope drops to zero (A); (ii) the force stabilizes to a constant force and the slope drops to zero (B) or (iii) the force continues to increase linearly but with a smaller slope until a second rupture event occurs (C) at which time the slope drops to zero. The drop in slope to zero when the force reaches a constant value indicates a membrane nanotube has formed. The measured loading rate: *r_f_*, the force to rupture: *f^R^* and lifetime: *Δt*^*R*^ are measured from the linear region of the force-time plot. Specifically, we determine the derivatives (j-slope^46^ 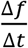 Δt: 75 or 100 ms) of the force-time plot where Δt is dictated by the signal-to-noise. Typically, the derivative (j-slope) before the onset of the tensile load undulates with magnitude of zero (0). Upon application of the load it increases and the time the first positive value is observed is defined as 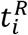 (Figure 2, Middle, A to C). It continues to increase to reach a near constant value and drops at time 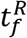 indicating a bond rupture event. The duration 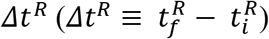 is defined as the lifetime of the bonds. After this event, the derivative either drops to a negative value and then climbs back to undulate around zero (Figure 2, Middle A and B) or drops to a lower positive constant value (Figure 2, Middle C). In the former case the force reaches a constant value after the rupture event, whereas in the latter the force continues to rise but with a lower slope until a second rupture event occurs (note: only the first rupture events are considered in this work). We define *f*^*R*^ as the force at 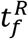. To verify the rupture event was not an artifact we calculate the j-slope and determine 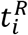 and 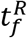 with larger Δ*t* (i.e. 125, 150, 175, 200 ms) and disregarded events that did not exhibit similar changes in the derivative. The measured loading rate *r_f_* is the slope of the linear fit to the region between 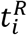 and 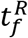 of the force-time plot and is 31.7, 32.1, 27.4 and 37.5 pN/s for examples shown in Figure 1D and Figure 2.

**Figure 2:**
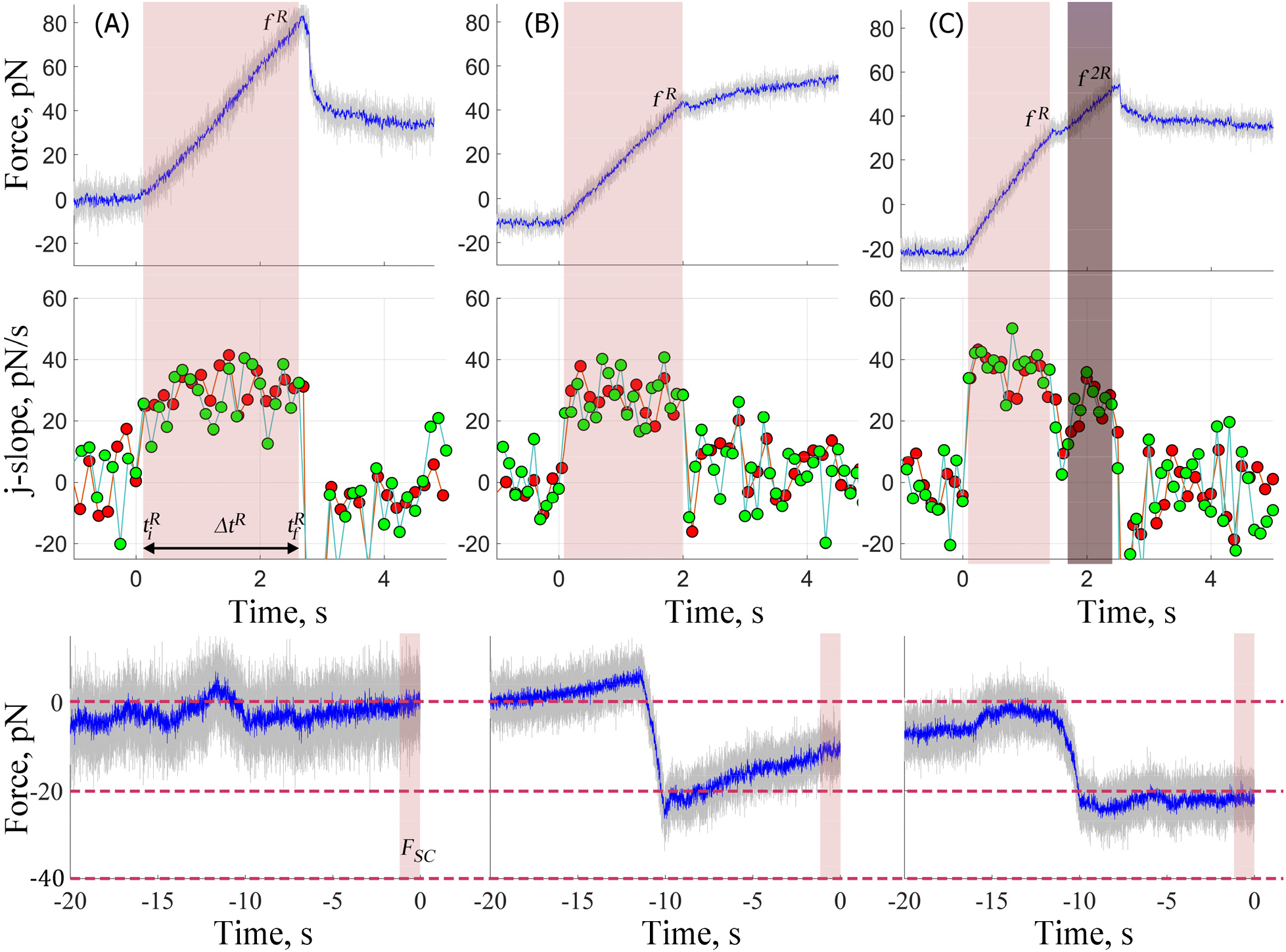
Determination of force and time to rupture. Force-time plots during the pull (**Top**) and respective j-slopes (**Middle**) and compressive force (**Bottom**) for three experiments. All experiments resemble one of the three typical cases. The force linearly increases with time until a rupture event occurring at time, 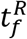 with force, *f^R^*. After the rupture event the force **A**: drops to a lower constant value having overshot the plateau force required to form the membrane nanotube 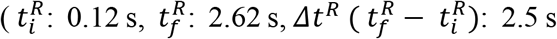 and *f^R^*: 82 pN and *r_f_*: 32.1 pN/s); or **B**: stabilizes at a constant value forming the membrane nanotube (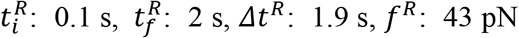 and *r_f_* 27.4 pN/s); or **C**: continues to rise linearly but at a lower slope and only reaches the constant value indicative of formation of a membrane nanotube after a second rupture event 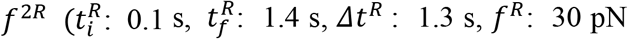 and *r_f_*: 37.5 pN/s). Bottom panels show the compressive force for the three experiments. The compressive force either stays close to 0 pN (**A**) or decreases below 0 pN from touch, until *t* = —10 s (**B & C**). For the next 10 s as the cell and bead adhere, it either remains stationary (**C**) or changes with a small slope (**B**) to reach a stationary value, *Fsc*. The magnitude of *Fsc* for three cases is 0, −11 and −22 pN. The green and red symbols represent j-slope calculated at 100 ms and 150 ms (**Middle**), see section 3.1 for details. Sampling frequencies are same as described in Figure 1.

### 2. Loading rate, *r_f_* depends upon the stationary compressive load (*F_SC_*)

We found the measured loading rate *r_f_* varies from 13 to 55 pN/s among experiments and that this range is partly explainable by the magnitude of the stationary compressive force upon approach, *F_SC_* (Figure 2, Bottom Panels). Shown in Figure 3A is a plot of *r_f_* as a function of *F_SC_; r_f_* decreases linearly with *F_SC_* by ∼-0.5 pN/s/pN to exhibit an unstressed *r_f_* of 26 pN/s, i.e.,

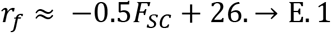

**Figure 3:**
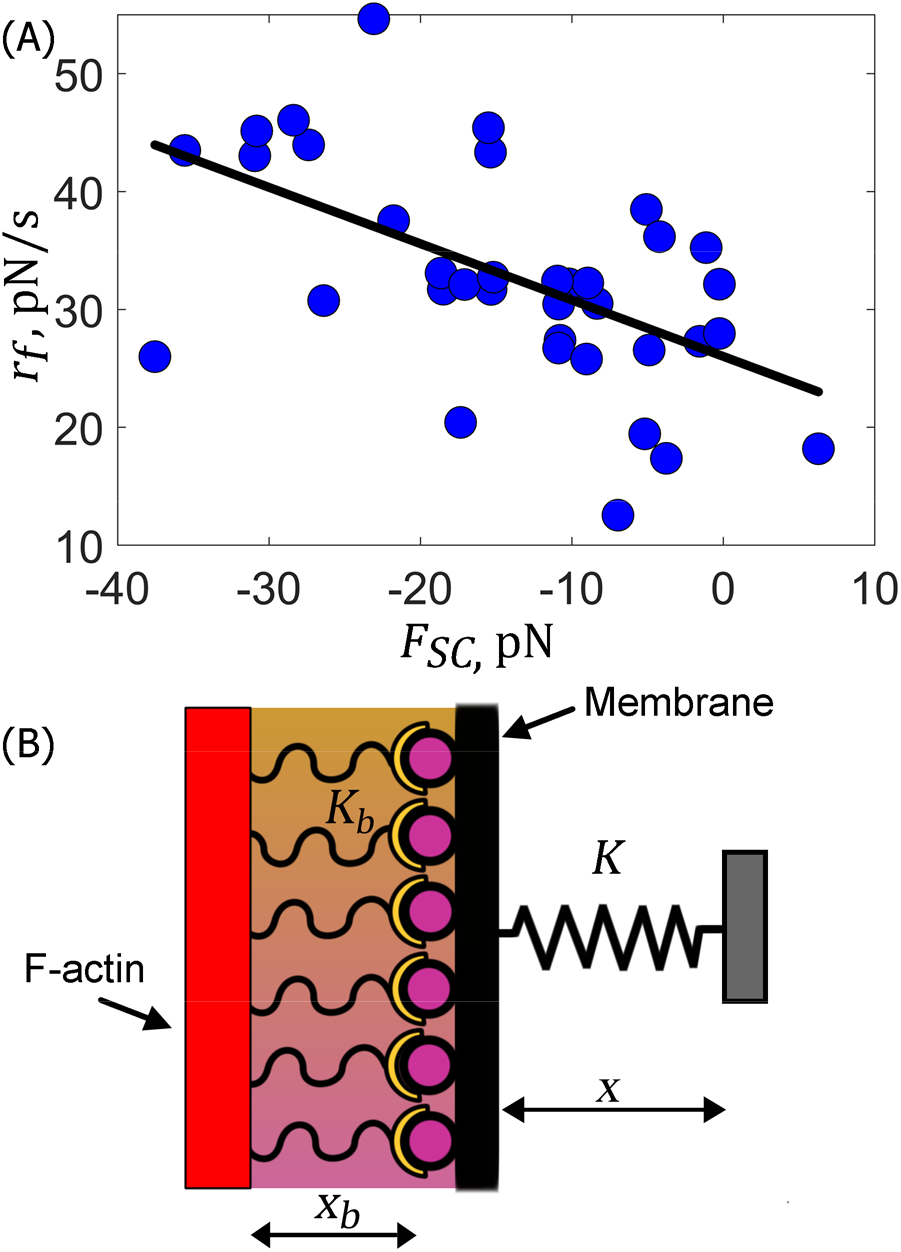
Loading rate, *r_y_* depends upon the stationary compressive load *(F*_*sc*_) for HN-31 cells. **A**:The *solid black* line is the best linear fit to the data; it has a slope of ∼-0.5 pN/s/pN (*P-value:* 5.4 × 10^-4^), an intercept of 26 pN/s (*P-value:* 1.9 × 10^-14^), and R^2^: 0.3. Each blue symbol represents a different experiment, number of experiments: 37. **B**: Schematic showing membrane cytoskeleton bonds with stiffness: *K*_*b*_ and extension: *x*_*b*_ in series with the transducer, i.e. optically trapped bead with stiffness: *K* and extension *x* when stationary.

This dependence of the loading rate on the compressive force can be explained if we assume the membrane-cytoskeleton bonds are in series with the probe and behave as Hookean springs^47^ with stiffness: *K*_*b*_ and *K* (Figure 3B). Force balance dictates that the force on the trapped bead, *Kx(t)* is equalized by the force on the bonds:

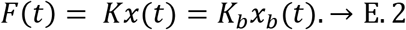

where *x_b_(t)* and *x(t)* are extensions of the bonds and the probe at time *t*. When the bonds are stretched at velocity *v*, geometry dictates that the total distance moved *vt* at time *t* is the sum of the displacement of the bead *x(t)* and extension of the membrane-cytoskeleton bonds *x_b_(t)*

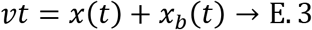

Solving E. 2 and E. 3 for *x*_*b*_*(t)* and equating the expressions, the force acting on the membrane-cytoskeleton bonds is then

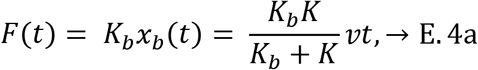

where the force acting on the bonds at 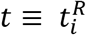 and 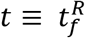 is

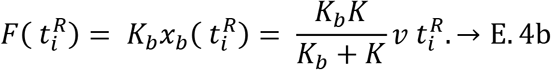

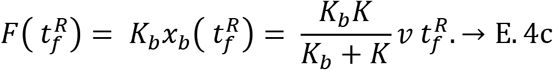

Subtracting E. 4b from E. 4c and rearranging noting that the measured loading rate, 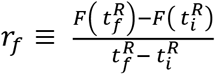 than

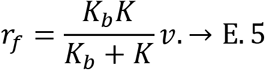

Substituting E.5 into E.1 we find

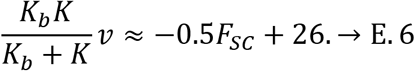

*K*_*b*_ is calculated from the ratio of the measured and nominal 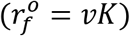 loading rates^25^ by rearranging E.5

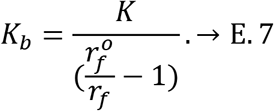

and found to vary from 0.02 to 0.39 pN/nm with mean: 0.162 ± 0.109 pN/nm. Because *K* and *v* are approximately constant among experiments then E.6 suggests *K*_*b*_ should increase as *F*_*sc*_ becomes more negative which is observed (Figure S1). This observation suggests the local regions of the cell react to applied compressive loads by increasing number of bonds or strengthening membrane-cytoskeleton bonds. Assuming a contact radius of 100 – 300 nm (see section 3.6) during compression, the mean local pressure on the membrane-cytoskeleton is ∼50 – 450 Pa, which is at least two-fold less than the compressive pressure found within native tumor microenvironment^48^ (∼770 Pa). This suggests the membrane-cytoskeleton bonds of HN-31 cells are mechanosensitive^49–51^ and can change their properties upon application of pressures or loads. This shows *F_sc_* should be restricted to minimize its effect on *r_f_*,; we note that *F*_*sc*_ was restricted to ≤ −10 pN in experiments conducted with neutrophils^25^, which would limit the increase in the loading rate to ≤ 20%, assuming both cells respond in a similar manner to the applied compressive load.

### 3. The stability of membrane-cytoskeleton bonds increases under smaller and decreases under larger tensile loading rates, *r_f_*

Data from HN-31 cells with comparable loading rates were binned (bin widths: ∼6-8 pN/s) and the means 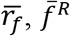 and 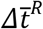 of *r_f_*, *f*^*R*^ and *Δt*^*R*^ determined. Figure 4A, (Left) shows the mean lifetime of the membrane-cytoskeleton bonds 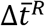 as a function of 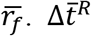 increases non-monotonically with 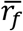 up to a threshold value of 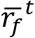 34.5 ± 2 pN/s. The maximum mean lifetime is reached at 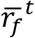 and is 2.2 s. As 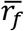 is increased beyond 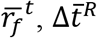 decreases non-monotonically. Figure 4B, (Left) shows 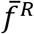 as a function of 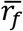. Like 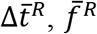 also increases with 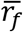 to reach 65 pN at 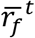, and then decreases with further increase of 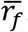. This shows that tensile loading rates up to the threshold stabilize membrane-cytoskeleton bonds, i.e. making them more difficult to separate while larger loading rates destabilize the same bonds.

The plot of 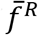 as a function of 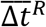 is shown in Figure 4C. The values increase linearly up to the threshold value of 65 pN (crosses) and then decreases linearly (open circles) exhibiting the same slope ∼34 pN/s. This linear relationship is predicted from E.4c because the force to rupture 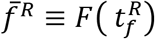 is proportional to 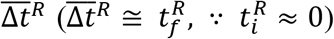. Indeed, the proportionality constant for this relation 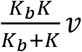 determined from the mean values of *K*_*b*_: 0.162 ± 0.109 pN/nm, *K:* 0.116 ± 0.011 pN/nm and *v:* 500 nm/s is ∼34 pN/s.

**Figure 4:**
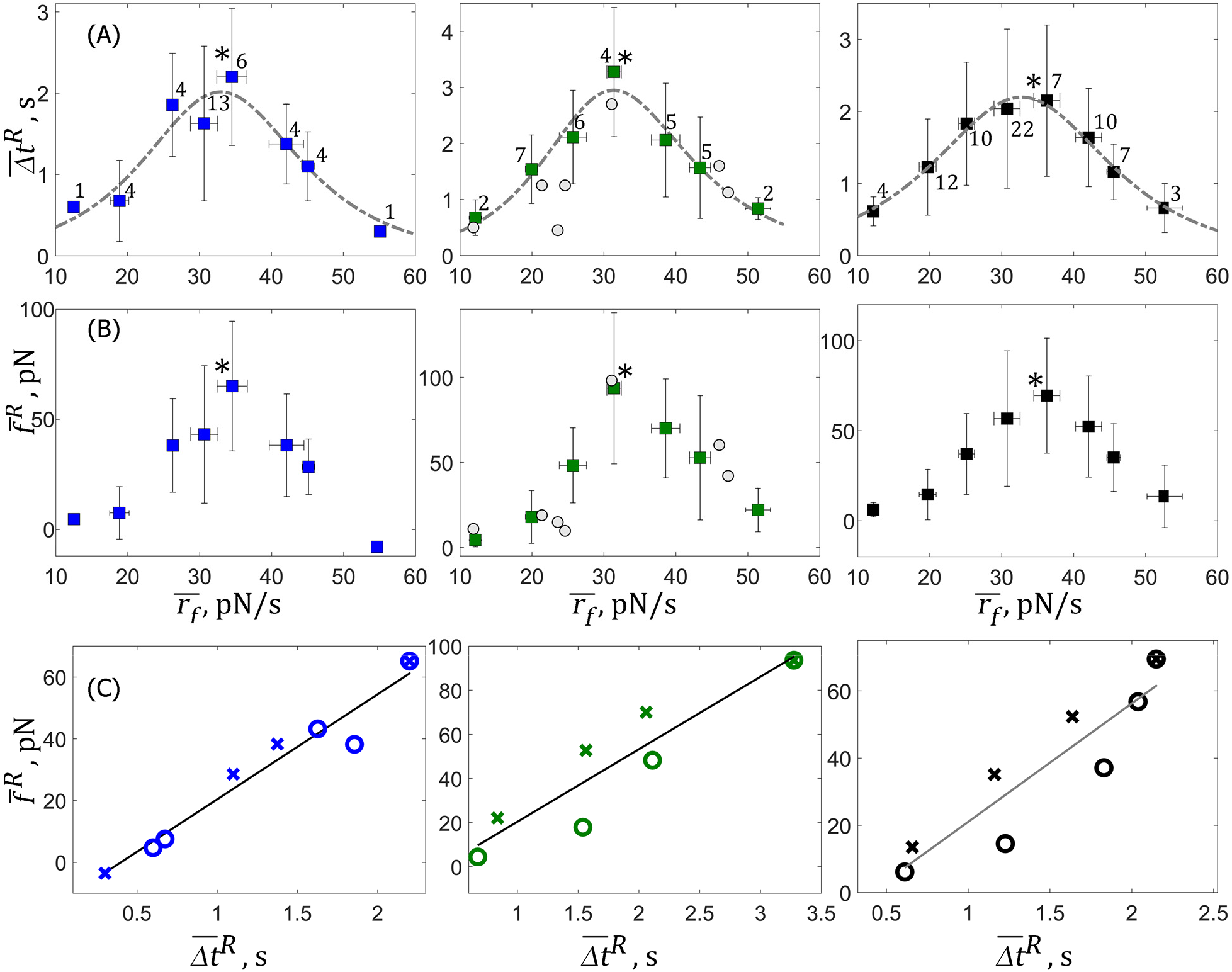
The stability of membrane-cytoskeleton bonds increases under smaller and decreases under larger tensile loading rates, *r_f_*. **A**: Mean lifetime of the bonds, 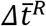 for untreated, (*blue*, **Left**) and Nocodazole-treated (*green*, **Middle**) cells as a function of loading rate. Individual experiments with DMSO-treated (*Grey circles*, Middle) cells are also shown. Threshold values 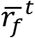 at maximum 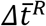 are noted by an asterisk. Shown in *black* (**Right**) is the mean response for cells from all three cohorts, i.e. includes untreated, Nocodazole-treated and DMSO-treated cells; maximum lifetime 2.2 s is reached at 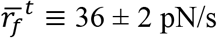. *Dashed grey lines* show the fits to E.16. Numbers indicate total number of different measurements at each loading rate. **B:** Mean force to rupture the bonds, 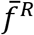; colors and symbols same as treatments described in **A.** The maximum 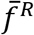 for the combined cohorts is 69.5 pN *(black*, **Right**). In **all panels** vertical bars represent standard deviations for 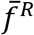 and 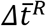 and horizontal bars represent standard deviations for 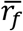 **C**: 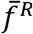 increases linearly with 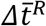 for all treatments. The open circles show small loading rates where 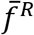 and 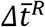 increase with 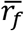 and the crosses represent larger loading rates where they decrease with 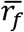. The *solid black* lines are the best linear fits with slopes 34 (R^2^: 0.95), 33 (R^2^: 0.84) and 35 pN/s (R^2^: 0.83) for the three panels respectively. Colors represent same treatments as found in A and B.

### 4. Membrane-cytoskeleton bonds depend upon F-actin integrity and active concentration of Rho-GTPases

F-actin is the predominant cytoskeletal protein in cells^52^ and is linked albeit not directly to the plasma membrane of cells. It is expected that drug treatments that inhibit polymerization of G-actin or promote depolymerization of F-actin should affect the membrane cytoskeleton bonds as reported^25^. To confirm this, we repeat the above experiments after treating the cells with an F-actin disrupting agent, Latrunculin A, as described in section 2.4. We first verified that Latrunculin A nearly depletes F-actin within HN-31 cells when incubated for times equivalent to the duration (20 to 60 min) of the experiment by imaging F-actin stained with Alexa 488 phalloidin. We found that F-actin was barely detectable (cf. untreated) when comparing the Latrunculin A-treated cells at the same (confocal) laser intensity as the untreated cells (data not shown).

Force-time curves with Latrunculin A-treated cells follow two typical patterns, an example for each (Top Panels) and its corresponding j-slope (Bottom Panels) are shown in Figure 5. Half (8/16) of the recordings, exhibit a short increase in force after which the force plateaus. For the example shown, the lifetime of the rise is 0.38 s with a *r_f_* of 23.0 pN/s (Figure 5A). The other half of the recordings exhibit a similar short increase but they are then followed by further non-monotonic rise in the force with time like the example shown in Figure 5B, where the lifetime of the initial rise is 0.6 s and *r_f_* is 30.7 pN/s. The first group exhibits similar characteristics to that described by Evans et al.,^25^ for Latrunculin A-treated neutrophils, i.e. the derivative of the force-time plot is decreasing with time to exhibit plastic-like behavior (cf. untreated). In their case, they suggest that this increase in force does not represent bond rupture but deformation of the membrane before it transitions to form a nanotube. They did not report examples as per our second group (Figure 5B). We suggest the reason for the slow and non-monotonic rise in the force before a nanotube is formed is due to presence of structures (e.g.: blebs or ruffles) that might diminish the membrane reservoir^53^. Blebs were observed at the base of the formed nanotube in some experiments.

**Figure 5:**
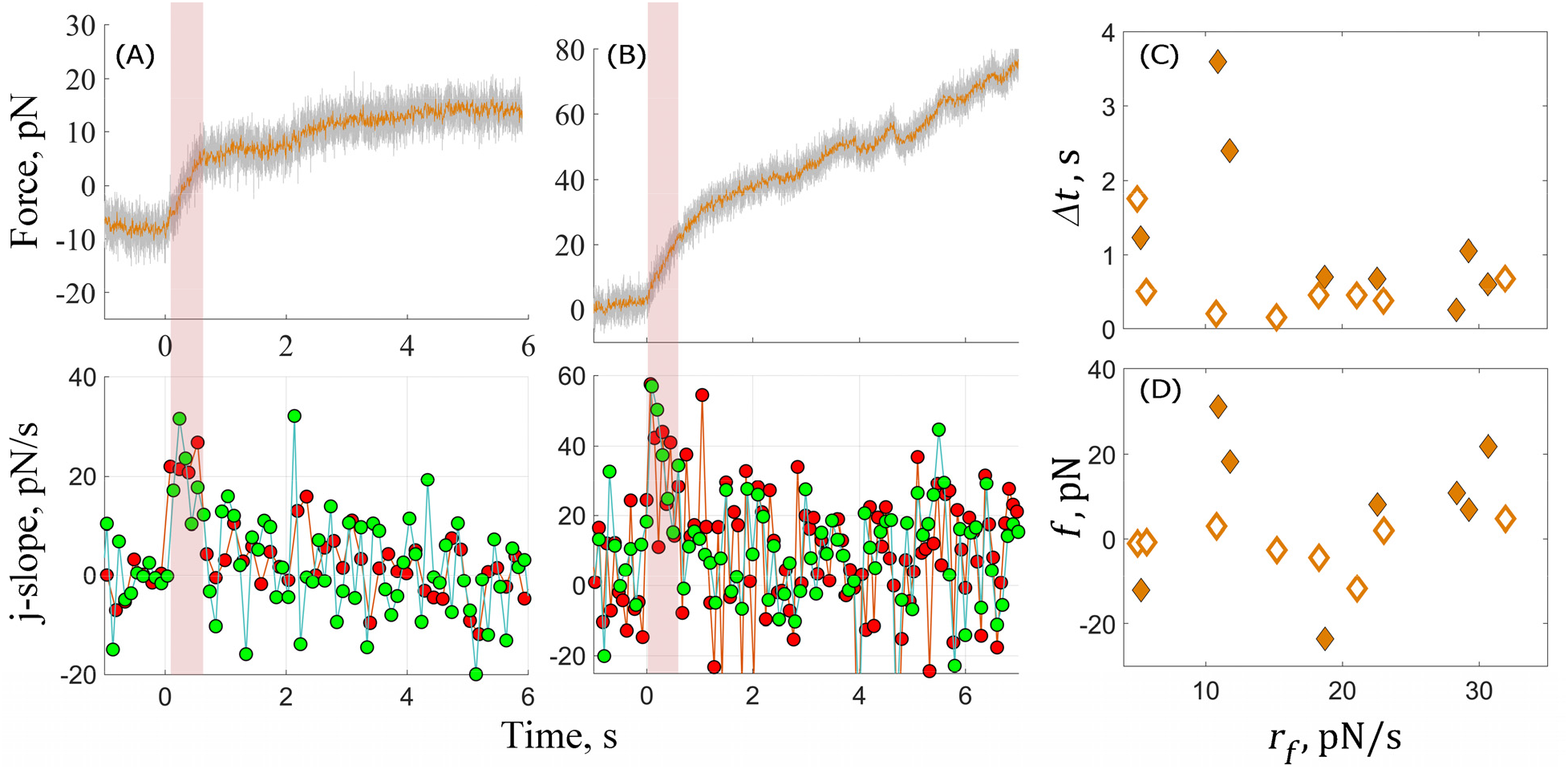
Mechanosensitivity at the membrane-cytoskeleton interface is no longer detected after Latrunculin A treatment. Force-time plots during the pull (**Top**) and respective j-slopes (**Bottom**) for two experiments are shown in **A** and **B**. All experiments resemble one of the two typical cases. The force increases with time for a short duration, *Δt* until a change in j-slope occurs at force, *f*, after which the force stabilizes at a constant value forming the membrane nanotube (*Δt:* 0.38 s, *f:* 1.89 pN and *r*: 22.1 pN/s, **A**); or continues to rise non-monotonically with time (*Δt:* 0.6 s, *f*: 21.75 pN and *r*: 30.7 pN/s, **B**). The green and red symbols represent j-slope calculated at 100 ms and 75 ms (**Bottom Panel in A**) or 100 and 150 ms (**Bottom Panel in B**). The sampling frequency of the force is 2000 (*grey*) and 200 Hz (*orange*) (**Top Panels, A & B**). *Δt* (**C**) and *f* (**D**) do not change with *r_f_* for both the first (open diamonds) and second (closed diamonds) groups.

The increase in force is slower compared to untreated cells and like Evans et al.^25^ we also suggest this increase does not represent bond rupture, but the deformation of membrane by the applied load. It is not surprising with such significant depletion of F-actin that the relationships observed with untreated cells (Figures 3A and Figure 4 Left Panels) are no longer observed; the cells are not mechanosensitive; the dependence of loading rate on compressive force is lost (Figure S2B). The data shows that the time (Figure 5C) and force (Figure 5D) are not dependent upon the tensile loading rate with both groups exhibiting mean *Δt:* 0.57 ± 0.50 s (median 0.45 s) and 1.2 ± 1.1 s (median: 0.7 s) with mean force −1.4 ± 5.1 pN (median −1 pN) and 7.6 ± 17.9 pN (median 9.3 pN). This data suggests an intact F-actin network is necessary for the cells to elicit mechanosensitive responses (Figures 2 and 3). We next examine candidates that enable linkages and affect the mechanosensitive response at the plasma membrane.

The actin cytoskeleton is linked to the plasma membrane via actin binding proteins (ABPs) either directly as observed for myristoylated alanine-rich protein kinase C substrate^54^ (MARCKS) and ezrin/radixin/moesin (ERM) proteins^55^, i.e. F-actin/ABP/Membrane or indirectly via membrane-linking proteins like Rho-GTPases. Examples of the latter are actin nucleators such as formins and actin-related protein 2/3 (Arp2/3) in a complex with its activator Wiskott-Aldrich Syndrome Protein (WASP), which can bind to membrane-bound Rho-GTPases to produce a linkage^56–58^ (e.g., F-actin/ABP/Rho-GTPase/membrane). Because Rho-GTPases regulate ABPs that bind F-actin to the membrane (e.g.: ERM proteins^55, 59^) and participate in complexes that directly link F-actin to the plasma membrane, we target them by use of *Clostridium difficile* Toxin B, a Rho-GTPase inhibitor. Toxin B is a monoglucosyltransferase that inactivates the Rho family GTPases (e.g. Rho, Rac and Cdc42) by catalyzing the monoglucosylation at either threonine 37 (Rho) or threonine 35 (Rac, Cdc42)^60^. These amino acids are part of the effector loop, i.e., the region on the Rho-GTPase that can bind proteins^61^ when the Rho-GTPase is in its active (GTP bound) state. Glucosylated Rho-GTPase, however, accumulates at the membrane in its GTP-bound form and is unable to cycle between the membrane and cytoplasm because it is unresponsive to GTP activating protein (GAP) and cannot be extracted by guanosine nucleotide dissociation inhibitor-1 (GDI-1)^62^.

We determined the efficacy of Toxin B on these cells by measuring the reduction in active RhoA-GTPase levels by a pull-down assay and found it reduced RhoA-GTPase activation by ∼27 ± 12 % after 2 hours of treatment and by ∼56 ± 22 % after 4 hours (Figure 6A & B). We then measured the reduction in F-actin levels at the membrane-cytoskeleton interface by staining F-actin with Alexa 488 phalloidin and compared the Toxin-B treated cells at the same (confocal) laser intensity as the untreated cells and observed a 25 % and 73% drop (from 156 ± 23 to 115 ± 33 and 42 ± 22) in F-actin levels after 2 and 4 hours of treatment (Figure 6C). We employed the 2 hour treatment for the rupture experiments.

**Figure 6:**
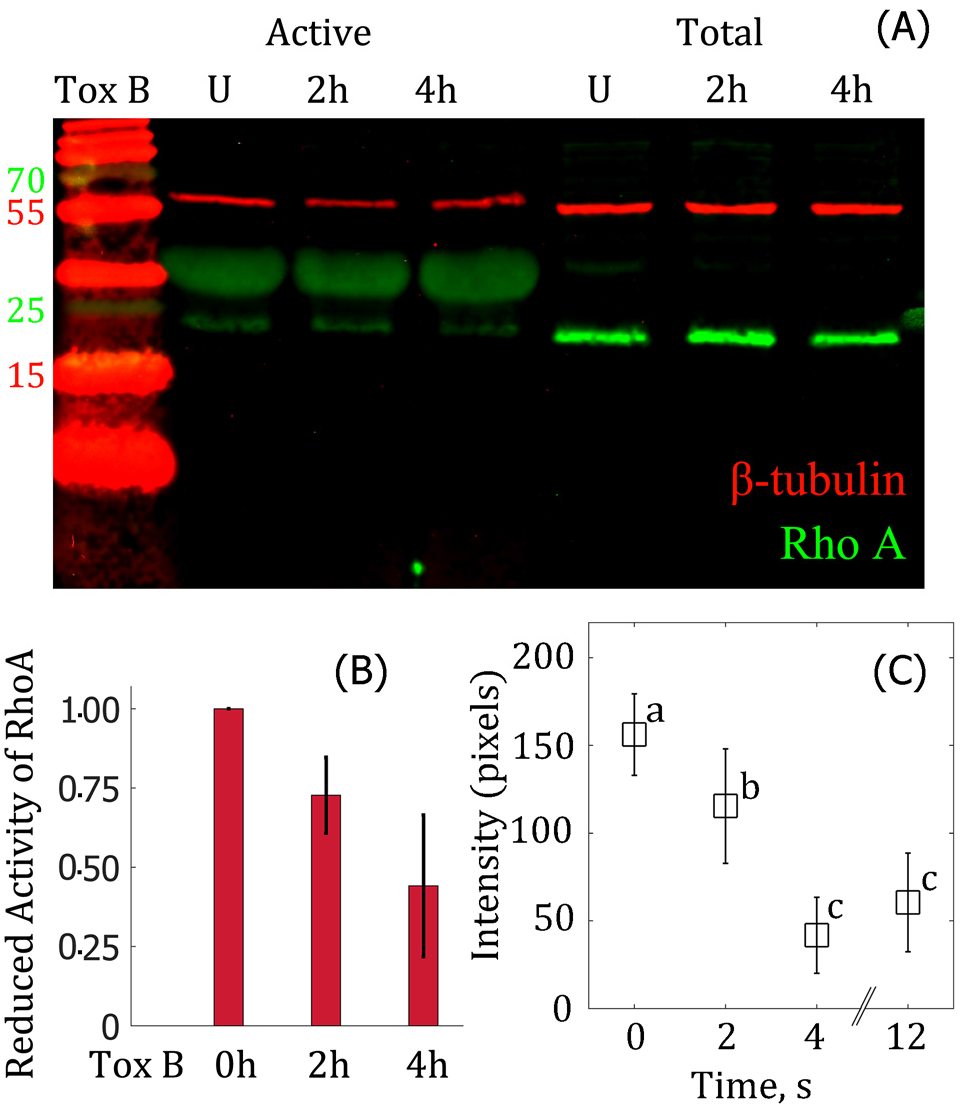
Toxin B diminishes active RhoA-GTPase concentration resulting in a reduction in F-actin levels at the membrane-cytoskeleton interface. **A**: Western blot where lanes represent first: ladder (molecular weights in kDa), second to fourth: active RhoA signal after pull down with Rhotekin-RBD beads and fifth to seventh: total RhoA. The symbols represent: U; untreated cells, 2 h and 4 h: 2 and 4 hour treatments with 200 ng/ml of Toxin B. RhoA is denoted by green (molecular weight: 21 kDa) and β-tubulin denoted by red (molecular weight: 55 kDa) is the loading control. **B**: Bar plot shows a reduction in active RhoA. Experiments repeated twice for each treatment. **C**: F-actin levels at the edge of the cell are reduced by ∼25 % after 2 hours and up to ∼70 % after 4 hours at the edge of the cell in the presence of Toxin B. The levels of F-actin were measured as the pixel intensity of the F-actin fluorescence (**Figure S3**). Different letters indicate statistical differences at *p-value* < 0.005 when assessed with the Tukey-Kramer test where number of cells per treatment is 10-14.

The force-time curves exhibit the same three patterns depicted for untreated cells in Figure 2 and we show two of them in Figure 7A and B. The first reaches a rupture force of 50.8 after 2.7 s at a loading rate of *r_f_* of 24.8 pN/s. The second example reaches a rupture force of 22.4 pN within 1 s, this was then followed by a second rupture event with a loading rate ∼ 4 pN/s lower than the first of 22.1 pN/s. Unlike experiments with untreated cells the stationary compressive force does not depend upon the loading rate (Figure S2C); for a similar number of rupture experiments we observe no relationship with this treatment. We find the lifetime 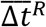 is also independent of loading rate to exhibit a mean value of 1.65 ± 0.55 s (Figure 7C, Bottom). The force to rupture 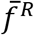 increases 4-fold with loading rate from 15 pN at 7.3 pN/s to 57.6 pN at 37.5 pN/s (Figure 7 D), and unlike with the untreated cells the mean lifetime is independent of the rupture force (Fig 7C, Top). Bonds that exhibit lifetimes independent of the load were observed between two cadherin monomers^19^. *K*_*b*_ for Toxin B-treated cells at 0.066 ± 0.056 pN/nm is on average 2.5-fold lower than *K*_*b*_ for untreated cells. This value is perhaps not surprising because activated Rho-GTPase is necessary to target F-actin to the membrane^56–58^ and our data (compare Figures 3 and 4 with Figures S2C and 7, and values of *K*_*b*_) suggests that it facilitates the formation of stiffer linkages that are capable of a mechanosensitive response. This assumes the measured decrease in active RhoA (cf. 27 ± 12 %, Figure 6) is comparable for the other Rho-GTPase proteins present in this cell^38^. The absence of the mechanosensitive response in Toxin B-treated cells leads us to suggest that the linkages are different than present in the untreated cells.

**Figure 7:**
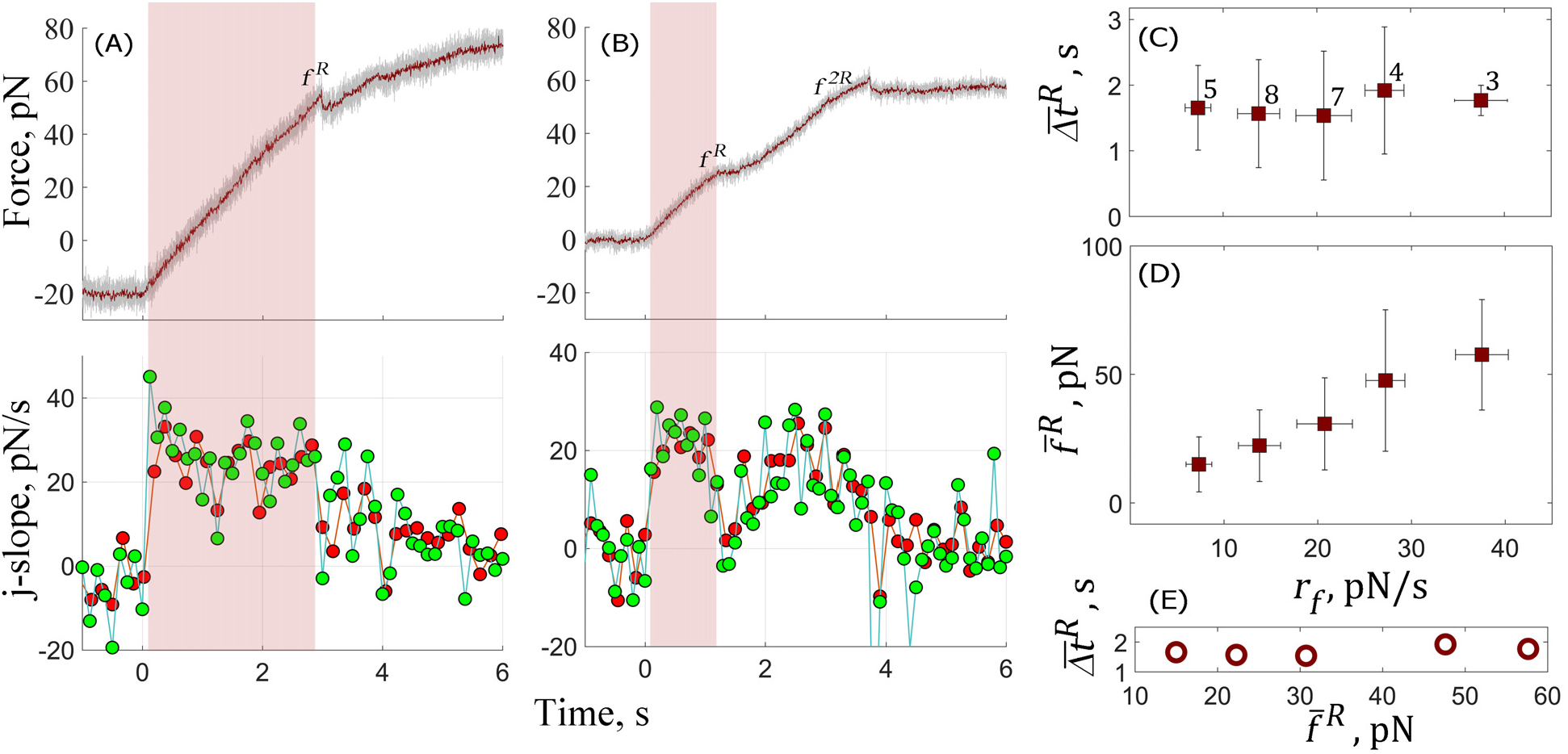
The lifetimes and force to rupture do not depend upon the tensile loading rate after cells are treated with Toxin B. All experiments for Toxin B-treated cells resemble one of the three typical cases for untreated cells shown in **Figure 2**. Force-time plots during the pull (**Top**) and respective j-slopes (**Bottom**) for two experiments are shown in **A** and **B**. The force linearly increases with time until a rupture event occurring at time, 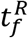 with force, *f^R^*. For the examples shown, after the rupture event the force stabilizes at a constant value forming the membrane nanotube (*Δt^R^*: 2.7 s, *f^R^*: 50.8 pN and *r_f_*: 24.8 pN/s, **A**); or continues to rise linearly but at a lower slope and only reaches the constant value indicative of formation of a membrane nanotube after a second rupture event *f*^*2R*^ (*Δt^R^*: 1 s, *f^R^*: 22.4 pN and *r_f_*: 22.1 pN/s, **B**). The green and red symbols indicate that j-slope was calculated at 100 ms and 150 ms (**Bottom Panels, A & B**). The sampling frequency of the force is 2000 (*grey*) and 200 Hz (*red*) (**Top Panels, A & B**). Mean lifetime of the bonds, 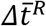 (**C**) and mean force to rupture the bonds, 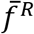 (**D**) for Toxin B treated cells. Open circles show 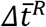 remains invariant as 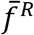 increases with 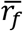 (**E**). Standard deviations for 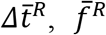 and 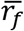 are shown and numbers (**C**) indicate total number of different measurements at each loading rate.

In contrast, disruption of tubulin polymerization by Nocodazole, seems to have no significant effect on the local stiffness determined to be 0.135 ± 0.09 pN/nm or their ability to respond to external compressive loads (Figure S2A) and tensile loading rates (Figure 3 Middle Panels). We first determined whether Nocodazole effectively disrupted the microtubule lattice in the HN-31 cell line by staining α/β-tubulin and imaging as described in section 2.4. We found that the lattice structure of the microtubules present in the untreated cells (Figure S4, Top Panels) and detected by the spatial Fast Fourier Transform was disrupted by Nocodazole treatment for the duration (20 to 60 min) of the rupture experiments. The cells retained their mechanosensitivity and modulated their membrane-cytoskeleton bonds in response to applied compressive loads (Figure S2A) and tensile loading rates. 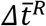 and 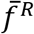 increase with the mean loading rate until 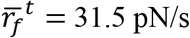 where they reach their maximum values of 3.3 s and 93.6 pN (Figure 4, Middle Panels A & B). The mean lifetimes and forces to rupture drop exponentially with further increase in 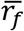. We found that cells when only DMSO was added exhibit similar behavior as that observed in untreated and Nocodazole-treated cells (Figure 4, Middle A & B, and Figure S2 A; *gray symbols*), confirming that the solvent is not overtly affecting the outcome. Figure 4 C shows the 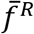 increases with the mean lifetime to exhibit a slope of ∼33 pN/s and resembles the untreated cells. Because we observe no significant difference among untreated, Nocodazole and DMSO treated we group them and recalculate the means of 75 rupture experiments. This is shown in the Right Panels of Figure 4. The 3 pN/s shift in 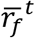 between the untreated (Figure 4A, Left) and Nocodazole-treated (Figure 4A, Middle) cells flattens the peak, observed when un-grouped (Figure 4A, Right) to result in a decreasing maximum sensitivity to the loading rate.

The lifetimes of the bonds show an unusual dependence on the tensile loading rate, they increase at low loading rates up to a threshold and shorten at higher loading rates. The former is akin to catch-like behavior while the latter mimics slip-like behavior of catch-slip bonds as observed in many single molecule experiments^14–17, 20–23^. Several groups have modelled catch-slip bonds. Pereverzev et al., proposed a two-pathway model^63^ where unbinding occurs via two alternative routes, catch and slip, where the applied force opposes the catch pathway while promoting the slip pathway. Later, they update their model^39^ to show that the force alters the conformational distributions of a complex and can be modelled with two pathways provided the transition between the two conformations is faster than bond dissociation. Two other models^40, 41^ also assume bond failure occurs from two bound-states and that the two states are in rapid equilibrium.

We use the model developed by Pereverzev et al. as it offers a direct comparison to a two-pathway model and identifies the origin of the catch-slip transition as a load-dependent conformational change. Because we apply a tensile loading rate (cf. force clamp) we rework their model to conform to our experimental conditions.

### 5. Catch-slip model for bond rupture upon application of a tensile loading rate

A kinetic scheme is shown in Figure 8. Consider that a cytoskeletal and membrane protein complex is present in either state 1 (B1) or state 2 (B2) and that it can transition between these states or dissociate via two separate pathways to produce two states (U1 and U2) of the unbound cytoskeletal protein and one of the unbound membrane protein which then equilibrate. (Note the kinetic scheme is also valid with one conformation of the unbound cytoskeletal protein and two conformations of the unbound membrane protein). We assume conformation B2 is more stable and the interaction with the membrane protein is stronger, hence it exhibits lower off-rates in absence of load (i.e., 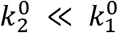). Therefore, unbinding occurs by first transitioning from state B2 to state B1 despite the equilibrium favoring state B2 (see E. 13, 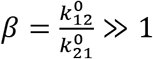). Hence in the absence of a tensile load the cytoskeletal-membrane-protein complex dissociates via state B1 to produce the separated membrane and separated cytoskeleton.

**Figure 8:**
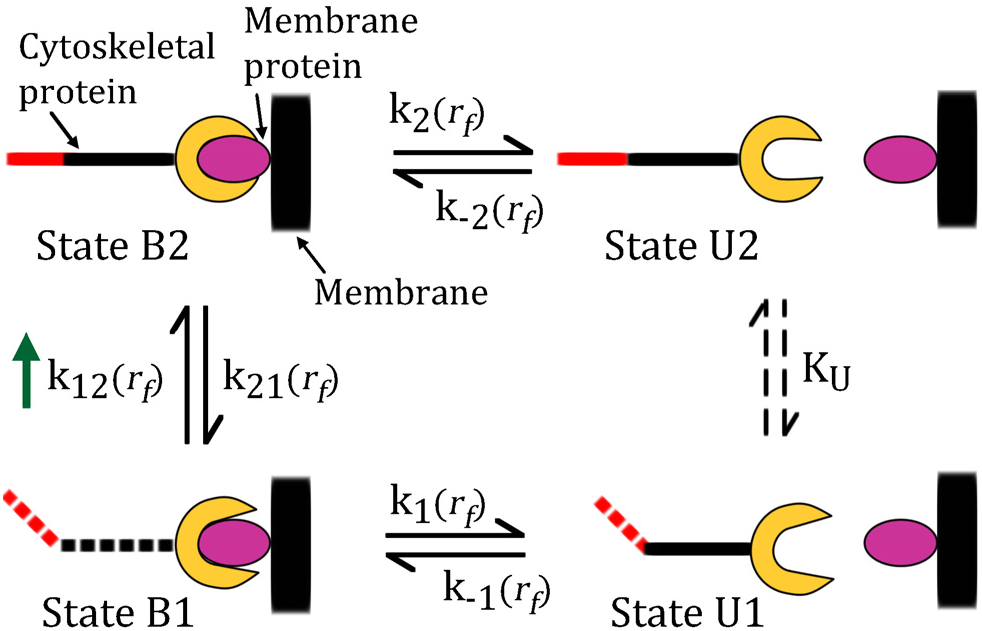
Kinetic scheme for membrane-cytoskeleton bond rupture via catch and slip pathways. A cartoon depicting each of the four different states is shown with the scheme. In this cartoon the complex is formed from a cytoskeletal protein (*yellow/black/red*) and membrane protein (*pink*) where we assume the cytoskeletal protein within the complex can exist in two conformations with state B2 (visualized as ‘extended’, *solid line*) more stable than B1 (visualized as ‘bent’, *dotted line*). When separated, the products equilibrate. Dissociation occurs via two pathways: (i) a catch-like pathway at no or low loading rates involving conformational change (B2 to B1) followed by dissociation (B1 to U1), i.e. *k*_21_ followed by *k*_1_ (ii) a slip-like dissociation at high loading rates (B2 to U2), i.e. *k*_2_ followed by equilibration between products. The rate constants for each step are shown and are: *k*_12_ and *k*_21_ for transitioning between B1 and B2; *k*_1_ and *k*_-1_ for dissociation and rebinding between B1 to U1; *k*_2_ and *k*_-2_ for dissociation and rebinding between B2 to U2. We assume the two unbound states U1 and U2 are in rapid equilibrium and denote this by a load-independent equilibrium constant (*K_U_*). All other rate constants are assumed to be load-dependent. Green arrow indicates *k*_12_ favors tensile load.

In our experiments, we apply a constant loading rate (*r_f_*). This tensile loading rate favors state B2 and increases the population of complexes in state B2 (relative to unloaded condition) while decreasing the number in state B1. At low loading rates (*r_f_* ≤ 34.5 pN/s) the complex continues to dissociate primarily via state B1, but due to reduction of population in state B1 with increase in *r_f_*, a catch-like decrease in off-rate (or increase in bond lifetime) is observed. At higher loading rates (*r_f_* ≥ 34.5 pN/s), the probability of finding the complex in state B1 is ≈ 0 and the bond starts dissociating from state B2 via a typical slip-like pathway. After bond dissociation via this pathway the unbound products reach equilibrium independent of the load. Based upon this scheme we now describe the model mathematically where we explicitly assume that the conformational transition between two bound states (B1 and B2) is much faster (µs time scale) than bond dissociation (≥ ms time scale). There is evidence for this for small proteins^64–66^ like Rho-GTPases (size ∼21 kDa) and G-actin (size ∼42 kDa). We also assume that dissociation is irreversible on the time scale, i.e. there is no rebinding as this should be small under a tensile loading rate. We first consider the equilibrium of the complexes between the two putative states B1 and B2.

Let *P_B1_(t,r_f_)* and *P_B2_(t,r_f_)* be the probability that the bound complex is either in states B1 and B2 then we can write

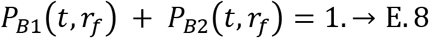

Because we assume the conformational transition between the two bound states is much faster than bond dissociation then at long times equivalent to time scale of experiment we write

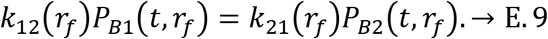

Rearranging and assuming the bonds are Bell-like^5, 47, 67^ then:

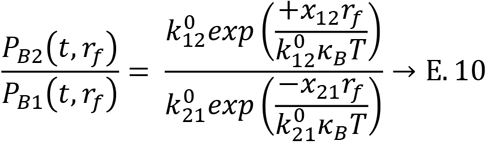

where 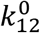 and 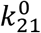 are rate constants in absence of force and *x*_12_ and *x*_21_ are the distances of the minima of states B1 and B2 from their respective activation barriers. The positive and negative signs indicate that while the applied tensile loading rate, *r_f_* promotes transition from state B1 to state B2, it obstructs transition from state B2 to state B1. Rearranging E.10 and simplifying we write

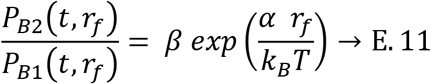

where, 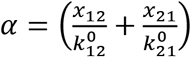 and 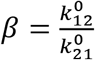 are constants associated with the conformational transition between the two states.

Making use of E.8 and E.11 the probability of finding the complex in either state is:

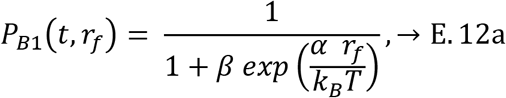

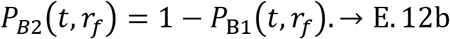

As we assume equilibrium favors state B2 then *β* ≫1 and E.12 becomes

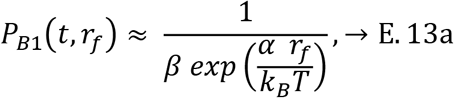

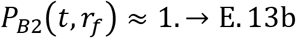

Assuming there is no rebinding the net dissociation rate of the bonds is the weighted average of the dissociation rates corresponding to each state and the probabilities of being in that state,

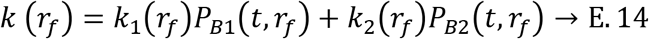

where 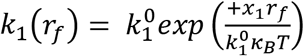 and 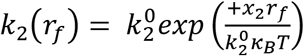 with unstressed rate constants 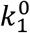 and 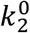 and barrier distances *x_1_* and *x*_2_. Note the tensile loading rate promotes both dissociation reactions hence barrier distances have the same sign. Substituting E.13 into E.14 then

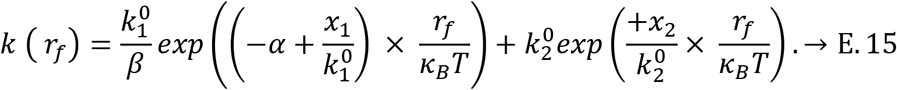

The lifetime (*Δt^R^*) of the bonds is then 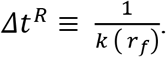

The six parameters obtained by fitting reciprocal of E. 15 to experimental 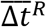 as a function of 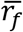 show that 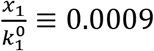 and 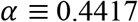, which implies 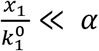, therefore we exclude 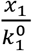, from E. 15 and re- fit the experimental data with only five unknown terms

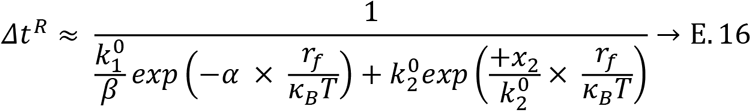

to find the parameters remain unchanged (*grey dashed line*, Figure 4, Table 1).

Now we re-write E.15

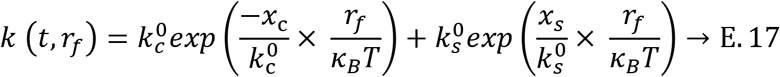

in terms of the catch-slip model^63^ modified to account for applied loading rate, where the unstressed rate constants for dissociation, and the energy barrier distances for the slip pathways are the same i.e., 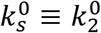 and 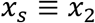 but for catch we find 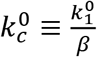 and 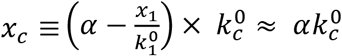 (because 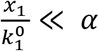) which we expand to find

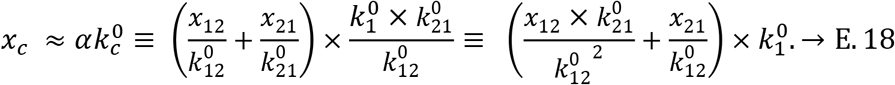

**Table 1:** Bond parameters determined by fitting experimental data to E. 16. and 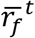 and 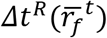 are determined from the maxima of the fit. Columns 5 and 6 are also slip parameters, catch parameters are calculated in columns 10 and 11. T: treatment, U: untreated, N: Nocodazole, and C: combined cohorts.

|  | Calculated parameters from the fit |  |  |  |  |  |  |  | Catch |  |
| --- | --- | --- | --- | --- | --- | --- | --- | --- | --- | --- |
| T | $\alpha$ | $\beta$ | $k_1^0$ | $k_2^0$ | $x_2$ | $\bar{r}_f^t$ | $\Delta t^R(\bar{r}_f^t)$ | $\varepsilon$ | $k_c^0$ | $x_c$ |
|  | nm.s | - | (1/s) | (1/s) | (nm) |  |  |  | (1/s) | (nm) |
| U | 0.4408 | 31.16 | 257.2 | 0.0099 | 0.0040 | 33.2 | 2 | 16.5 | 8.25 | 3.6 |
| N | 0.5345 | 31.02 | 262.6 | 0.0104 | 0.0040 | 31.2 | 2.9 | 24.6 | 8.47 | 4.5 |
| C | 0.3724 | 49.13 | 222.6 | 0.0104 | 0.0040 | 31.8 | 2.2 | 10.0 | 4.53 | 1.7 |

This shows the *x*_*c*_ only depends upon the barrier distances for the conformational transition between B1 and B2.

We determine these parameters by comparing E.16 with E.17 for the catch-like pathway and the slip-like pathway (Table 1, columns 5, 6, 10 and 11) for untreated HN-31 cells (Table 1, row 1). We find that treatment with Nocodazole (Table 1, row 2) does not significantly affect calculated parameters of the fit for the membrane-cytoskeleton bonds. We do find the efficiency^63^, *ε* of a catch bond is higher (24.6) for Nocodazole-treated cells compared with untreated cells at 16.5. It is the ratio of lifetime at the threshold loading rate 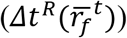 and unstressed lifetime 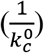. The reason for the difference is because threshold lifetime is ∼1 s shorter for untreated compared to Nocodazole-treated cells. For the grouped dataset (Figure 3, Right A), the shape of the bell widens and peak flattens and this is reflected by the apparent lower efficiency of 10. More data are necessary to establish whether there is a subtle difference between treatments or the combined cohort better represents the response to force.

In the absence of an applied loading rate (*r_f_* = 0) we calculate from E. 12, 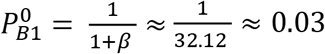 and 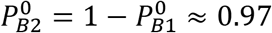. As predicted, the dominant state is B2 but dissociation rate from this state is small at 0.0099 s^-1^ (Table 1, column 5). In contrast, although probability for the complex to be in state B1 is low at 0.03 the dissociation rate is determined to be 257 s^-1^ (Table 1, column 4). Therefore, as expected dissociation rate from state B1 is greater at 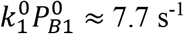 than from state B2 at 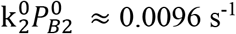.

The off-rates and energy barrier distances predicted for the conformational change by this kinetic scheme are of the same order as others have found^39^. The reported catch barrier distance was 26.8 Å and 22.5 Å for P-selectin/PSGL-1 and FimH/mannose, respectively, while we found it to be 36.4 Å and 17 Å for membrane-cytoskeleton bonds of untreated cells and the combined cohort, respectively. This distance reflects the folding or unfolding distance between the two conformational states and is expected to be greater than dissociation distances. The barrier distance calculated for slip pathway is 0.04 Å, which is too small to represent dissociation of a single bond (expected to be around 1 Å) and suggests we may have a cluster of bonds. If we assume *N* bonds are sharing the load uniformly the net dissociation rate *k* (*r_f_*) is defined by E. 14 but with 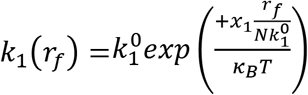 and 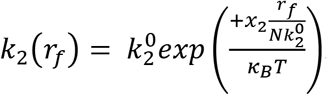. Substituting them into E. 14, simplifying and excluding 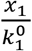 (because 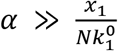) as shown above, we would find the first term in the 2^nd^ exponent of the denominator for the modified E.16 is 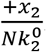. This slip-barrier distance determined with the multi-bond model is *N* times larger than the value calculated with the single bond model as shown for Bell bonds^68^.

Indeed one theoretical study suggests a biphasic loading rate time relationship should only be detectable when a cluster of *N* bonds ruptures fractionally under uniform loading conditions^69^ and hence would be difficult to experimentally detect. We note that our biphasic relationship was observed at low loading rates (< 60 pN/s) which allowed *N* bonds to rupture fractionally (Figure 2C, Figure 7B).

### 6. Nanotube formation force, plateau force and evidence that F-actin grows within the lumen of the formed nanotubes

The onset of nanotube formation is determined from the last membrane-cytoskeleton bond rupture event^25^; following it the force then stabilizes to a plateau value (Figures 1D and 2). We divide the formation force into two groups: overshoots where the force either goes past and then drops (Figure 2A and 2C) or non-overshoots where the force stabilizes (Figure 1D and Figure 2B and Figure 7) to a constant plateau value, *fº*. We find the median (mean) and standard deviation of the force to form a single nanotube for the overshoots and non-overshoots is 54 (62) ± 15 (8) and 39 (45) ± 25 (6) pN respectively (number of experiments are in parenthesis). The formation force did not vary significantly when the cells were treated with Nocodazole (overshoot: 68 (67) ± 24 (7) pN, no-overshoot: 42 (39) ± 10 (4) pN) and Toxin B (overshoot: 60 (77) ± 34 (5) pN, no-overshoot: 35 (34) ± 12 (7) pN). The force reaches *fº* with median (mean) and standard deviation of 37 (44) ± 18 pN and the nanotube extends to a length of 10 µm (Figure 1D). Multiple nanotubes form in some experiments where the plateau force increased as discrete multiples of the single nanotube *fº* (∼35 pN) (Figure 9A). The plateau force did not vary significantly when the cells were incubated with Nocodazole or Toxin B. There was no difference between formation 21 (19) ± 13 (8) pN and plateau value 19 (19) ± 11 pN when F-actin was disrupted with Latrunculin A, both were about half the magnitude observed with untreated cells (group I only since group II does not plateau, Figure 5). Surprisingly, we observe no significant difference in the nanotube formation and plateau forces in Toxin B-treated in comparison with untreated cells. This contrasts with the difference we observed when we examined the time course of the rupture force (compare Figures 4 versus 7) in untreated versus Toxin B-treated cells. The measurements reflect different processes. The time course of the rupture force reflects characteristics of bond breaking in response to the tensile loading rate, i.e. the mechanosensitivity of the cell whereas the plateau force indicates there is a readily available membrane reservoir to extend the nanotube^70^. The magnitude of the plateau force depends upon the pulling speed and approaches a minimum value as *v* approaches zero^71–73^. The plateau forces reported for cells vary from 20 to 65 pN for different *v* (range: 0.076 to 3 µm/s) and different temperatures (room, 24 °C versus physiological, 37 ºC). When comparing our results with experiments conducted at same *v, fº* observed for nanotubes formed from NIH3T3 fibroblasts^34^ (39 to 42 pN, 0.076-0.908 µm/s) is of the same magnitude, whereas it is 10 pN more for neutrophils^73^ (51 pN) and 10 pN less for Human Embryonic Kidney (HEK) cells^72^ (∼30 pN). It is 15 to 20 pN less than our measurements for Human Brain Tumor (HB) cells^74^ (19 pN, 3 µm/s), Chinese Hamster Ovary (CHO) cells^74^ (23.2 pN, 3 µm/s), neurons^75^ (∼15 pN, 1 µm/s) despite the faster pulling rate employed in these studies. The plateau force for astrocytes^75^ and glioblastoma cells^75^ (32 pN, 1.0 µm/s) is of the same order as our measurements while that found for microglia^75^ (∼60 pN, 1 µm/s) and macrophages^75^ (∼65 pN, 1 µm/s), is 10-20 pN more than HN-31 cells albeit *v* was twice as fast. Others have also found the plateau force is reduced when F-actin is disrupted as reported for HEK cells^72^ (24 pN), NIH3T3 fibroblasts^34^ (23 pN), HB cells^74^ (13 pN) and neutrophils^73^ (22.5 pN).

**Figure 9:**
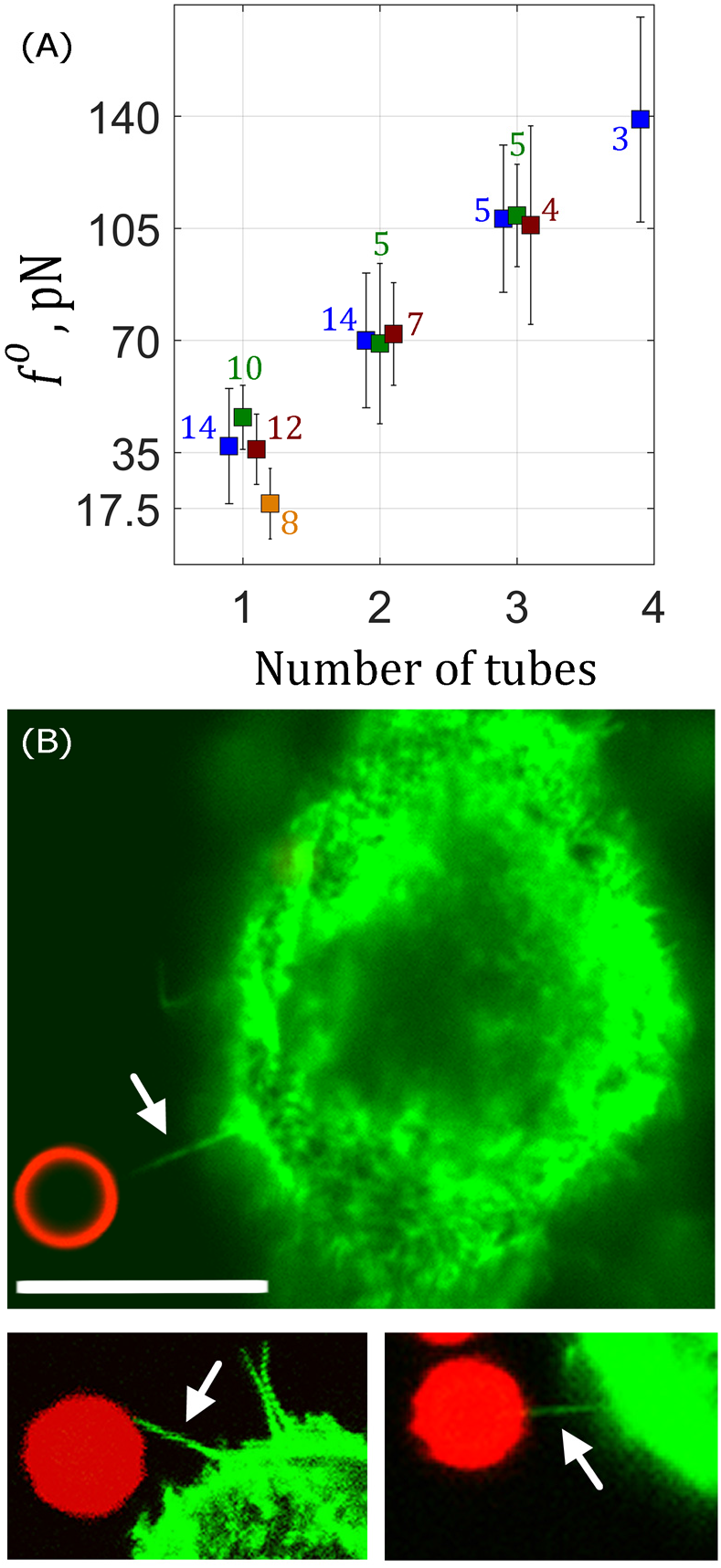
F-actin is localized in the formed nanotube. **A**: The plateau force *fº* is the same for untreated (blue), Nocodazole (green) or Toxin B (red) treated cells and lower for Latrunculin A (orange) treated cells. *fº* increases in quantiles with the number of nanotubes. The medians (means) and standard deviations are 37 (44) ± 18, 70 (74) ± 21, 108 (107) ± 23, and 139 (137) ± 32 (untreated); 46 (46) ± 10, 69 (77) ± 25, 109 (109) ± 16 (Nocodazole), and 36 (39) ± 11, 72 (71) ± 16, 106 (103) ± 31 (Toxin B) pN for one (1), two (2) and three (3) nanotubes. The numbers indicate the total number of independent measurements. **B**: Images of HN-31 cells stained for F-actin with Alexa 488 phalloidin (green fluorescence) show that a thin nanotube containing F-actin (white arrows) is present between the cell and bead. The bead is visible in red fluorescence. The scale bar is 10 µm.

Koster et al.^76^ showed for membrane vesicles that the force barrier for the formation of a membrane nanotube increased linearly with the area the force is exerted on. Pontes et al.^34^ demonstrated this with NIH3T3 fibroblasts by showing the ratio 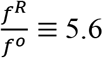 was dependent on the ratio between radius of formed nanotube and radius of contact area between bead and cell (coined the patch radius). We find the ratio is smaller at 1-1.5 which suggests that if E.16 of Pontes et al. is valid, the contact patch area is two to three-fold greater than nanotube radius, i.e., for nanotube radius of 50 – 100 nm^34, 74, 77^ the patch radius is predicted to be 100 – 300 nm. We do find as shown for many different cell lines^34, 75^ that membrane nanotubes formed from HN-31 cells contain F-actin (Figure 9B). The samples were fixed < 1 minute after the onset of the tensile load (Figure 1) and this represent a time scale comparable to our force experiments. We previously suggested that membrane nanotubes are F-actin filled to explain sawtooths (transient rise in force followed by decay) that rode above a stationary value^46^. In that report we hypothesized the sawtooth transients arose from depolymerization and polymerization of F-actin that pulled and pushed the membrane. We observed similar sawtooths in HN-31 cells^26^ and they have also been reported in neurons^78^ and HeLa cells^79^. We also detect inverse sawtooths, i.e. a transient decay in force followed by a rise^80^ and suggest that F-actin present in the nanotube (Figure 9B) polymerizes at the tip of the nanotube to push the membrane forward resulting in a decay in the force. We detect inverse sawtooths especially once the applied load is removed but in two experiments also during nanotube extension, between 10 and 20 seconds after *Pull (Start)*. The imaging of F-actin and the detection of the transients suggest that HN-31 cells like the fibroblasts, neurons and supporting cells readily fill the lumen of the formed nanotubes with F-actin. We note HN-31 when cultured *in vitro* exhibit considerable number of filopodia (∼150) and our results show that small tensile loads may be sufficient to initiate their formation.

## 4. Conclusions

1. We observe a biphasic relationship where the membrane-cytoskeleton bonds increase their lifetimes as loading rates increase to a maximum, after which they exhibit shorter lifetimes with further increase in loading rates. This behavior resembles catch-slip bonds demonstrated with purified protein systems *in vitro* and can be explained by a load-dependent conformational change that precedes dissociation of membrane-cytoskeleton bonds by either catch or slip pathways. Our experiments were conducted by rupturing the bonds at relatively low tensile loading rates unlike other studies where the load is clamped at a constant value until bond rupture. Performing experiments at lower loading rates, and under force clamp will help validate our methodology.
2. We demonstrate that the cells lose their ability to respond to mechanical cues when concentration of active Rho-GTPases is reduced. We speculate the mechanosensitive linkages involve actin binding proteins where Rho-GTPases either directly participate or regulate the linkages. Experiments performed upon inhibition of actin binding proteins (e.g.: formins, WASP, ERM proteins, MARCKS) and interacting motor proteins (e.g.: myosin II, 1b) also capable of forming catch bonds, will help to elucidate the molecular origin of the bond.
3. Our findings suggest local molecular ensembles at the edge of HN-31 cells are sensitive to mechanical stimuli, the bonds increase their lifetimes in response to low tensile loading rates and stiffen in response to small compressive loads. This stimulus initiates cytoskeleton remodeling observed as growth of F-actin within the lumen of membrane nanotubes that form post-bond rupture.

## 6. Acknowledgement

Research supported by NIH grants S10 RR027549-01, R21CA152779, and RO1DC00354 and by the Alliance for Nano-health 1W81XWH-10-2-0125. We appreciate the advice of Dr. Jeffrey N. Myers to use the HN-31 cell line in our studies, and his generous gift of the cell line. We would like to thank Ameeta A. Patel for providing assistance and training in the optimum conditions needed to culture this cell line and Dr. Michelle L. Seymour and Haiying Liu for assistance and training with the pull-downs assay and Western blot techniques.

---

**Figure S1:**
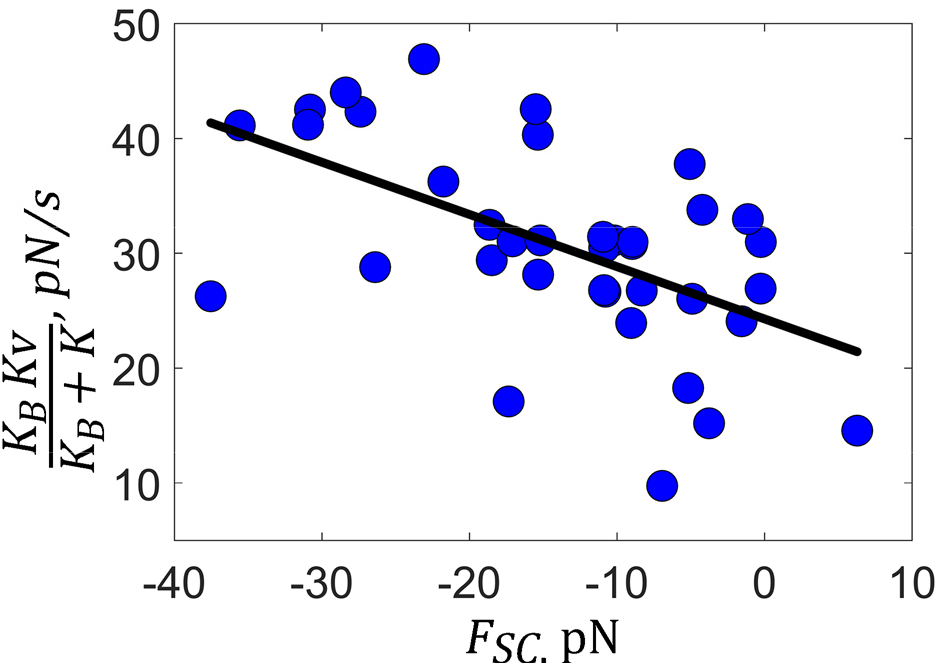
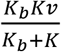 decreases linearly with *F*_*sc*_ for HN-31 cells (*blue*). The best-fit line (*solid, black*) has a slope of ∼-0.45 pN/s/pN (*P-value*: 3.6 × 10^-4^), an intercept of 24 pN/s (*P-value*: 4.0 × 10^-14^), and R^2^: 0.3. Each blue symbol represents a different experiment, Number: 37.

**Figure S2:**
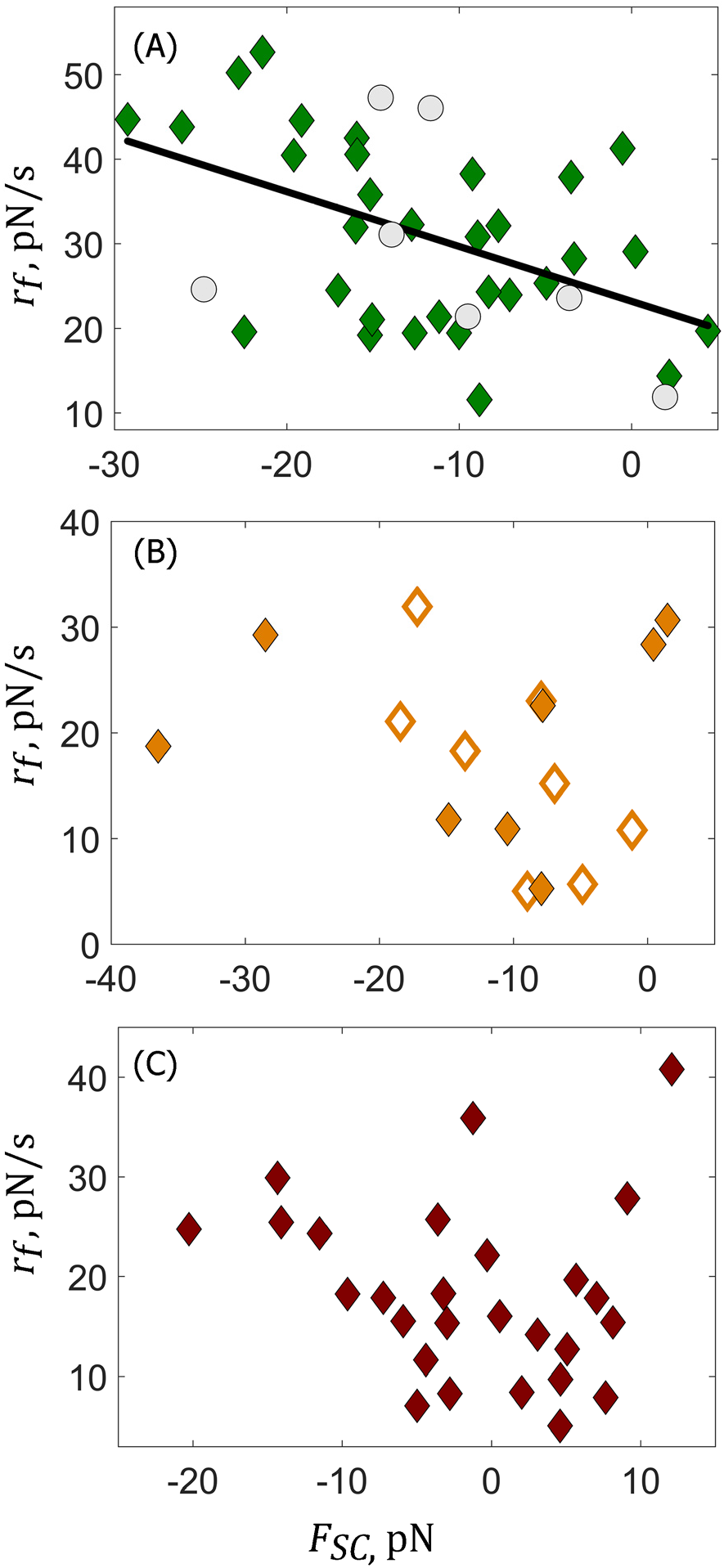
**A**: *r_f_* retains its linear dependence with *F_SC_* in cells treated with Nocodazole (*green*). The best fit line (*solid black*, R^2^: 0.25) has a slope of ∼-0.7 pN/s/pN (*P-value*: 4.2 × 10^-3^) and an intercept of 23 pN/s (*P-value*: 4.8 × 10^-8^). Grey symbols show the experiments performed with cells with only the solvent DMSO added. Latrunculin A *(Orange, open symbols: first group and closed symbols: second group*, **B**) and Toxin B (*red*, **C**) treated cells do not exhibit linear relationship (*p-values* of the slopes are 0.56 and 0.47). Each symbol represents a different experiment.

**Figure S3:**
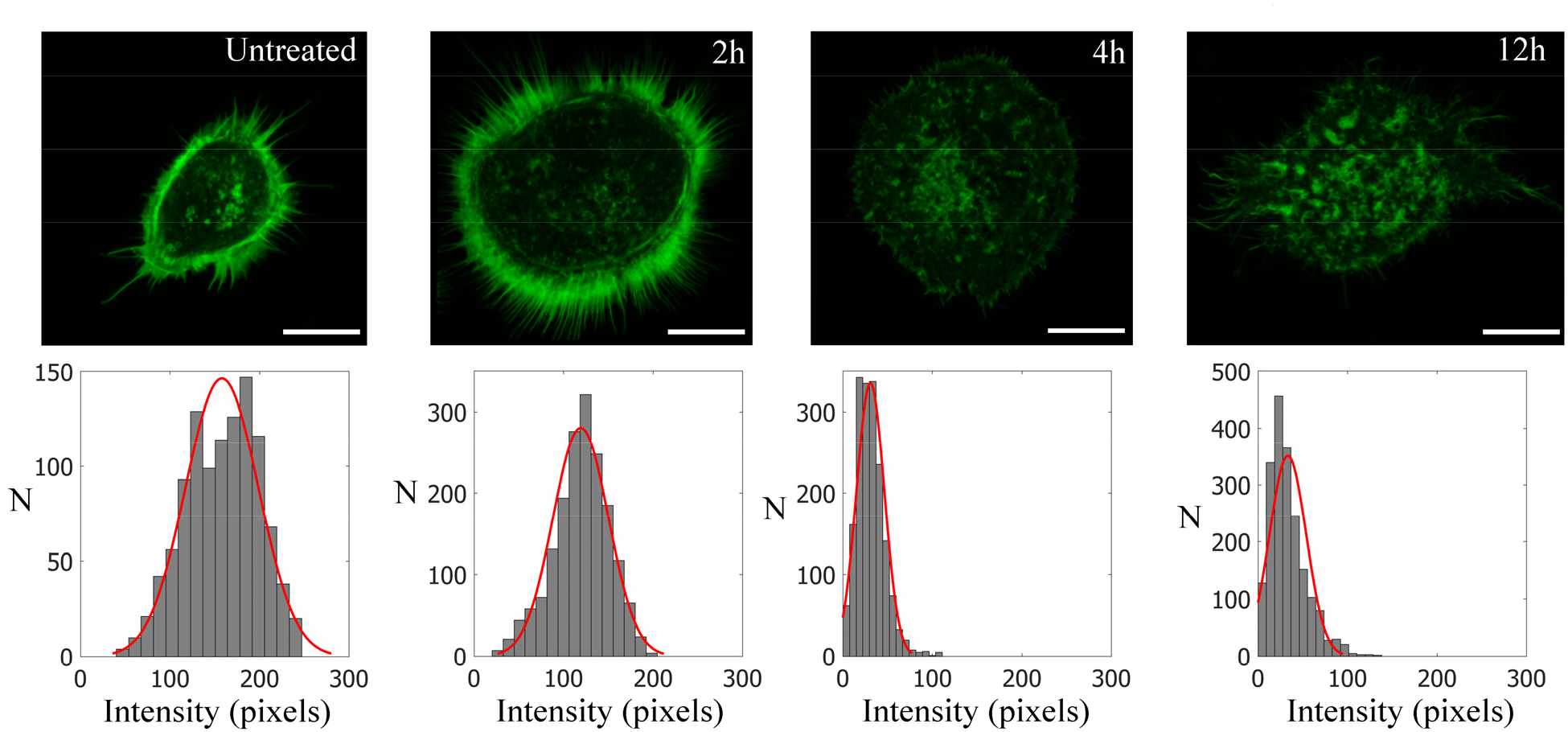
Method to determine the pixel intensity of F-actin fluorescence at the membrane cytoskeleton interface. **Top Panels** show confocal images for untreated and cells treated with Toxin B for 2, 4 and 12 hours. F-actin levels at the membrane-cytoskeleton interface are determined from the intensities (8-bit: 0 to 255) of the labelled F-actin (green) at the boundary of each cell in an image (the cell boundary was hand drawn and the intensity at the boundary was determined with MATLAB). **Bottom Panels** show histograms of intensities obtained for the corresponding images. The red line is the fit to a normal distribution with mean intensities (**Left to Right**) 150, 119.3, 30.8 and 33.4. Each image is the average of 16 slices of the cell at different heights starting at the bottom of the dish.

**Figure S4:**
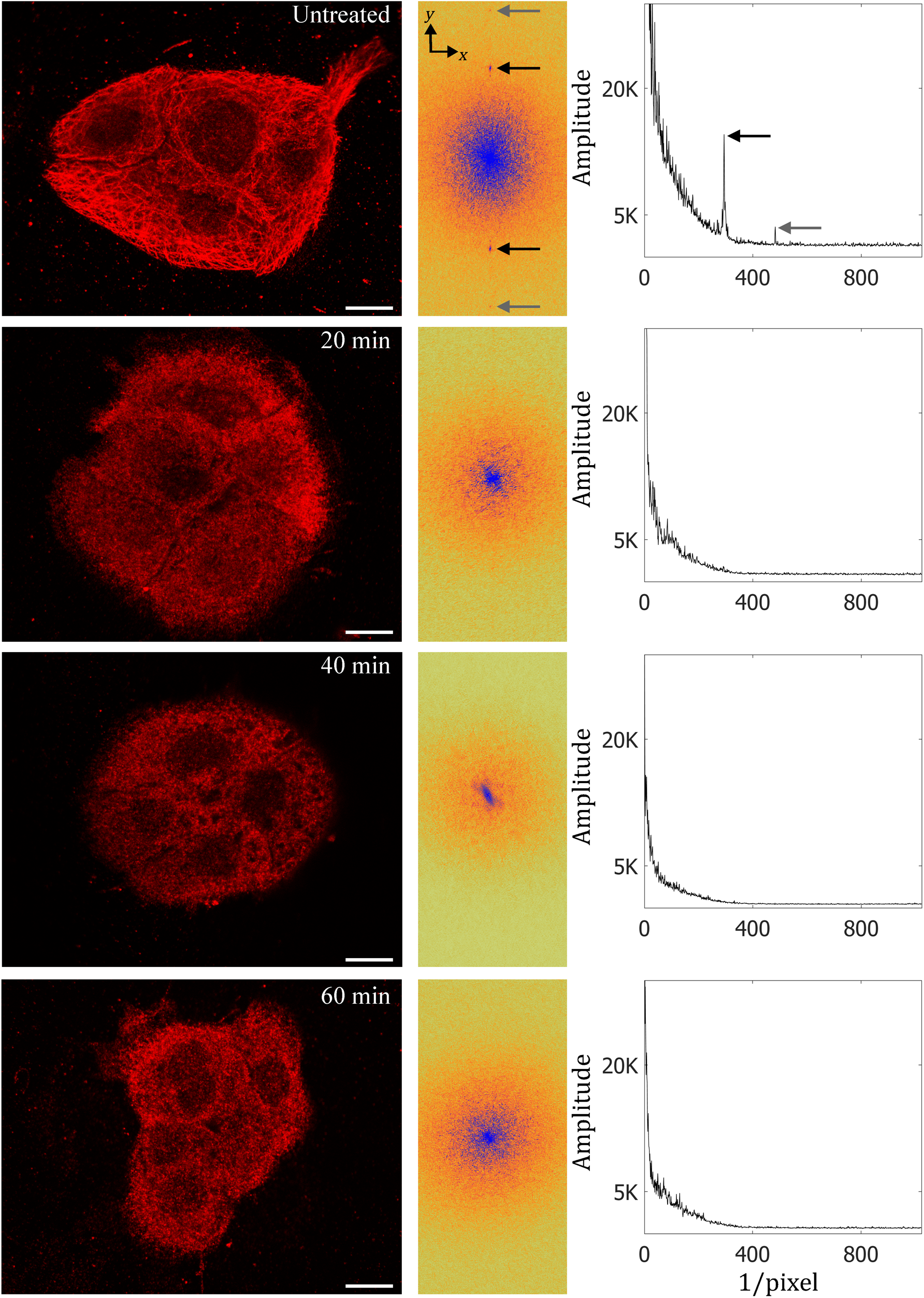
Nocodazole disrupts the microtubule lattice in HN-31 cells. **Left panels** show confocal images of HN-31 cancer cells with β-tubulin labeled in red for untreated and after 20, 40 and 60 minutes of treatment with Nocodazole. **Center panels** show the spatial Fast Fourier Transforms (FFT) of the corresponding confocal images. Harmonics are detected only in untreated cells, and are found from the center position along the *y*-axis, in both positive and negative directions. The arrows show the harmonics; the black arrow shows the first and the grey the second harmonic. **Right Panels** show a slice of the FFT from the center position along the *y*-axis in the negative direction. Harmonics when present in the **Center Panels** are visible as peaks. The peaks are only detected in untreated cells.

**Table S1:** Table of abbreviations.

| Parameter | Description | Units |
| --- | --- | --- |
| $X, Y$ | Positional information of the QPD in the $x$ & $y$ directions | V |
| $N$ | Number of bonds | - |
| $\Sigma$ | Sum, intensity information of the QPD | V |
| $m_{QPDx}$ | Linear slope between bead displacement and normalized voltage | V/V/nm |
| $x, y$ | Positional vector of the bead in Cartesian coordinates | nm, nm |
| $K$ | Stiffness of the trap | pN/nm |
| $r_m, \theta$ | Positional vector of the bead in polar coordinates | nm, radians |
| $v$ | Velocity of the piezo stage | $\mu\text{m/s}$ or $\text{nm/s}$ |
| $t$ | Time | s |
| $F$ | Force | pN |
| $\Delta x$ | Displacement of the bead in the $x$ direction in nm in time $\Delta t$ | nm |
| $f^R, \bar{f}^R$ | Tensile force to rupture the bonds & mean tensile force to rupture bonds | pN |
| $\Delta F / \Delta t$ | The local derivative (j-slope) | pN/s |
| $t_i^R$ | The time j-slope becomes positive during application of tensile loading rate | s |
| $t_f^R$ | The time of a bond rupture event | s |
| $\Delta t^R, \Delta \bar{t}^R$ | Lifetime of the bonds & mean lifetime of the bonds | s |
| $F_{SC}$ | Stationary compressive load | pN |
| $K_b$ | Local elastic stiffness of the bonds | pN/nm |
| $x_b$ | Bond extension | nm |
| $r_f, \bar{r}_f, r_f^o$ | Measured loading rate, mean measured loading rate & nominal loading rate | pN/s |
| $x(t)$ | Displacement of bead at time, $t$ | nm |
| $x_b(t)$ | Extension of bonds at time, $t$ | nm |
| $F(t)$ | Force at time, $t$ | pN |
| $f_b$ | Force scale, $f_b = \frac{\kappa_B T}{x_b}$, where $\kappa_B$: Boltzmann constant & T: Temperature | pN |
| $k_1(r_f), k_2(r_f)$ | Rate constants for bond dissociation from states B1 & B2 under $r_f$ | 1/s |
| $k_1^0, k_2^0$ | Off-rates for bond dissociation from states B1 & B2 under no load | 1/s |
| $x_1, x_2$ | Widths of the barrier associated with each well | nm |
| $k_{12}(r_f), k_{21}(r_f)$ | Rate constants for transition from B1 to B2, & B2 to B1, under $r_f$ | 1/s |
| $k_{12}^0, k_{21}^0$ | Rate constants for transition from B1 to B2, & B2 to B1, under no load | 1/s |
| $x_{12}, x_{21}$ | Distances of the minima of state B1 & B2 from the activation barrier | nm |
| $\alpha$ | Conformational factor | nm.s |
| $\beta$ | Equilibrium constant between states B1 & B2 | - |
| $P_{B1}(t, r_f), P_{B1}^0$ | Probability the bound complex is in B1, at $t$ and under $r_f$, & under no load | - |
| $P_{B2}(t, r_f), P_{B2}^0$ | Probability the bound complex is in B2, at $t$ and under $r_f$, & under no load | - |
| $k_c^0, k_s^0$ | Rate constant for dissociation via the catch and slip pathways | 1/s |
| $x_c, x_s$ | Catch and slip barrier distances | nm |
| $r_f^t$ | Threshold loading rate at which $\Delta \bar{t}^R$ & $\bar{f}^R$ are maximum | pN/s |
| $\Delta t^R(\bar{r}_f^t)$ | Maximum lifetime obtained at threshold loading rate | s |
| $\varepsilon$ | Efficiency of the catch bond | - |
| $f^o$ | Plateau force | pN |

